# Restoring oxidative phosphorylation enhances osteogenesis in mitochondrial DNA translation defective human bone marrow stromal cells

**DOI:** 10.1101/2024.12.24.629993

**Authors:** Paula Fernandez-Guerra, Pernille Kirkegaard Kjær, Simone Karlsson Terp, Jesper S. Thomsen, Blanca I. Aldana, Herma Renkema, Jan Smeitink, Per H. Andersen, Johan Palmfeldt, Kent Søe, Thomas L. Andersen, Moustapha Kassem, Morten Frost, Anja L. Frederiksen

## Abstract

Bone formation is critical to maintain bone integrity. Here, we studied the importance of intact energy metabolism for bone formation in humans. The skeletal impact of impaired oxidative phosphorylation (OXPHOS) was investigated in adult individuals with genetically defective mitochondrial DNA translation (m.3243A>G). Although impaired mitochondrial ATP production in m.3243A>G human bone marrow stromal cells (hBMSC) was compensated by increased glycolytic ATP production (unchanged net ATP production), both *in vitro* osteoblast differentiation and *in vivo* ectopic bone formation were decreased. The impaired OXPHOS was associated with mitochondrial stress and disruption of the pro-osteogenic transcriptional program characteristic of hBMSC. Supporting OXPHOS pharmacologically in hBMSC restored mitochondrial ATP production, their transcriptional program and metabolism, leading to upregulation of osteogenic genes and restoration of bone formation capacity. These findings demonstrate a mitochondrial regulation mechanism of the osteogenic capacity of hBMSCs and identify OXPHOS as a potential target for increasing bone formation.

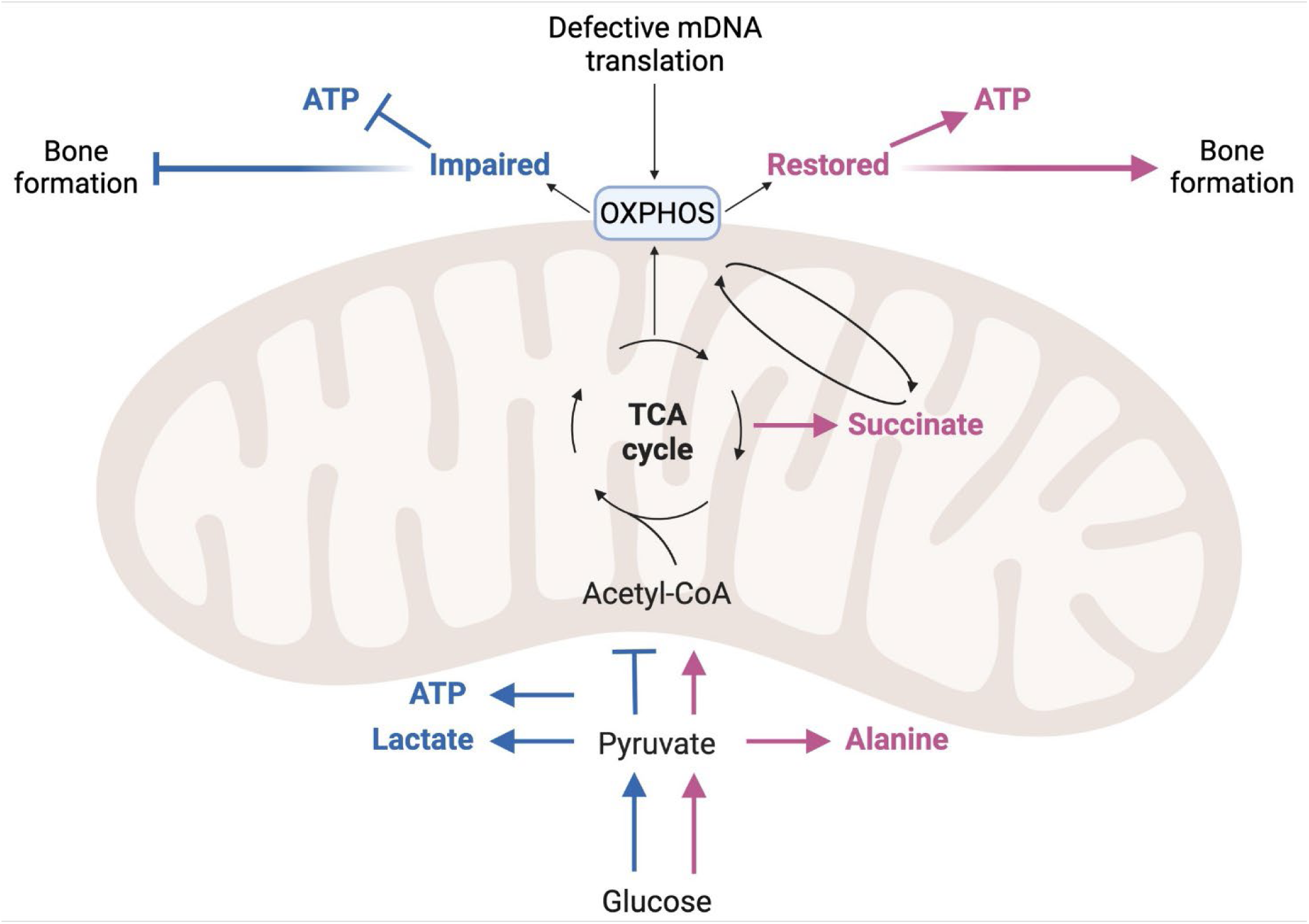

## Introduction

Bone is a dynamic tissue that is remodelled throughout life to maintain bone mass equilibrium and mechanical competencies. Bone remodelling, the continuous process of bone degradation by bone-resorbing cells – osteoclasts – followed by the formation of bone matrix by bone-forming cells – osteoblasts^1^ is a highly energy-demanding process^2,3^. The primary energy source for bone tissue is glucose, and accordingly, skeletal sites with substantial bone remodelling have high glucose uptake, as demonstrated by radio-tracer studies in adult women and men^4,5^. Cellular energy, in the form of adenosine triphosphate (ATP), is generated via mitochondrial oxidative phosphorylation (OXPHOS) and cytoplasmatic glycolysis^5^. The significance of sufficient ATP for bone has been studied in osteoprogenitor cells (human bone marrow stromal cells, hBMSC), which prefer glycolysis for proliferation and self-renewal in agreement with somatic stem cells from other tissues^5^. In contrast, data on osteoblasts differentiated from hBMSCs is less uniform. During osteogenic differentiation, most studies show OXPHOS activation while glycolysis remains unchanged^5^. Additionally, mouse osteoprogenitors exhibit metabolic flexibility by shifting from OXPHOS to glycolysis after pharmacological inhibition of the respiratory chain complex III and vice versa after lactate dehydrogenase (LDH) inhibition, where increased OXPHOS activity exerted a bone anabolic effect^6,7^. The importance of OXPHOS to skeletal integrity is further supported by the finding of low bone mass, bone deformities, and fragility fractures in mice with impaired ATP supply caused by OXPHOS deficiency in respiratory chain complex I, III, IV, and V^8–10^. Mitochondria are multifunctional organelles, and in addition to decreased ATP production, impaired OXPHOS function affects other mitochondrial functions, including the production of tricarboxylic acid (TCA) cycle intermediates, regeneration of NAD^+^, and regulation of mitochondrial stress responses (MSR)^11–14^. The extent to which these processes influence bone homeostasis in humans remains to be established.

While *in vitro* and animal studies demonstrate the significance of intact OXPHOS for osteoprogenitor differentiation and the development of normal bone structure, clinical data linking mitochondrial dysfunction – specifically impaired OXPHOS – to skeletal integrity are limited. Increased bone fragility was reported in an epidemiological study of subjects with inherited mitochondrial disorders^15^, suggesting an association between mitochondrial dysfunction and compromised bone properties. In addition, we previously reported thinner cortical bone and lower bone mineral density (BMD) in adult women and men carrying mitochondrial DNA (mDNA) pathogenic variant m.3243A>G in the tRNA^Leu(UUR)^ gene (*MT-TL1)* that leads to defective translation of mDNA-encoded proteins and consequently to OXPHOS impairment^16^.

Collectively, pre-clinical and observational studies suggest that impaired OXPHOS activity compromises bone cell differentiation and function, potentially impairing skeletal integrity. However, the role of mitochondria in bone and its clinical significance remains understudied. Here, we used osteoprogenitors (hBMSCs) from symptomatic and asymptomatic m.3243A>G carriers to investigate the impact of mitochondrial dysfunction on human bone cells. We show how a highly glycolytic bioenergetic profile in osteoprogenitors reduces their osteogenic potential by regulating their transcriptional program. Furthermore, the pharmacological enhancement of OXPHOS activity in the osteoprogenitors restored their pro-osteogenic transcriptional program and subsequent osteogenic capacity. Overall, these findings extend the knowledge of the effect of OXPHOS on hBMSCs, osteoblasts and bone formation as well as propose OXPHOS as a potential therapeutic target for improving bone formation and supporting skeletal health.

## Results

### hBMSC from m.3243A>G carriers present a transcriptional, proteomic, and metabolic profile of impaired OXPHOS with a compensatory highly glycolytic profile

We have previously shown that individuals carrying the mitochondrial DNA (mDNA) pathogenic variant m.3243A>G have lower bone mineral density (BMD), thinner cortices, and lower estimated bone strength^16^, which indicate that mitochondrial function influences the maintenance of skeletal integrity. The pathogenic variant m.3243A>G is located in the *MT-TL1* gene that encodes for the tRNA^Leu(UUR)^, leading to impaired translation of the OXPHOS proteins encoded in mDNA. To further detail the effect of impaired OXPHOS on bone, we did a comprehensive assessment of the impact of m.3243A>G on osteoprogenitors (hBMSCs), bone cell function, *in vivo* bone formation, and clinical measurements, including dual-energy X-ray absorptiometry (DXA). We recruited 10 adult subjects carrying the pathogenic variant m.3243A>G; tRNA^Leu(UUR)^ defined as carriers - including individuals with diabetes and/or myopathy as well as asymptomatic individuals – and 10 healthy controls closely matched on age, sex, and body mass index (BMI) **(Figure 1A)**. There was no difference between groups in plasma levels of major regulators of bone homeostasis, such as vitamin D (25OHD), Ca^2+^, parathyroid hormone (PTH), thyroid hormone (TSH), and liver and kidney function **(Table S1)**. As previously reported^16^, resting plasma lactate levels were significantly higher in m.3243A>G carriers (Fold-change (FC) = 1.58, *p* < 0.01) **(Figure 1B).** The increased plasma lactate results from increased glycolysis to compensate for the impaired ATP production from OXPHOS^17,18^.

**Figure 1.**
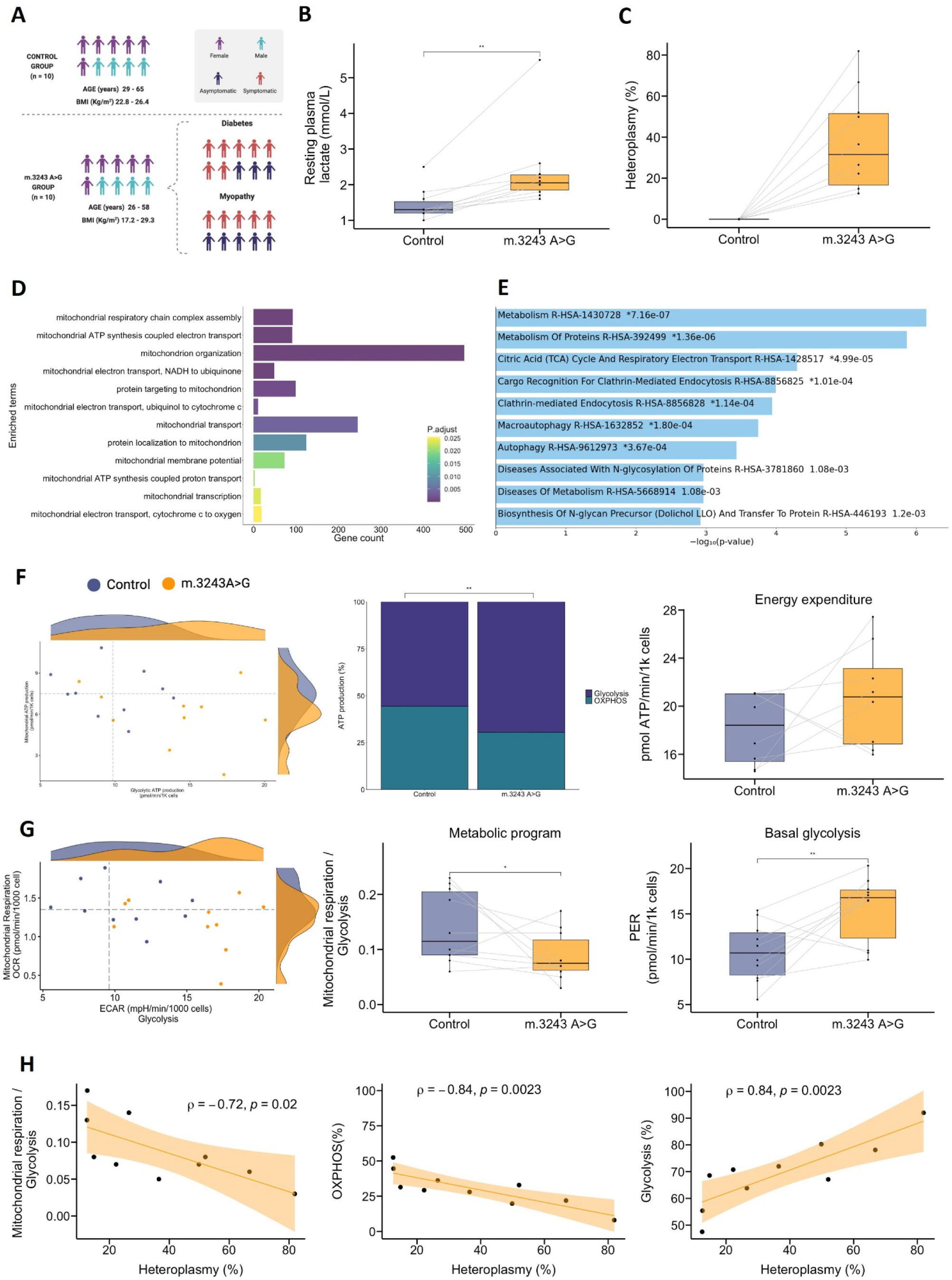
hBMSC from m.3243A>G carriers show a more glycolytic profile. **A)** Study cohort with inherited defective mitochondrial DNA translation (m.3243A>G mDNA) and healthy individuals matched by age-, sex-, and BMI. **B)** Resting plasma lactate. **C)** Heteroplasmy of hBMSC analysed by ddPCR. **D)** Enrichment analysis related to mitochondrial function of differentially expressed genes of hBMSC from subjects with m.3243A>G compared to controls using EnrichR with GO biological process terms as database. The enriched terms show the number of genes in each GO term and the adjusted *p*-value of the GO term. **E)** Enrichment analysis of differentially expressed proteins of hBMSC from m.3243A>G carriers compared to controls using EnrichR with Reactome as database. The enriched terms are shown as negative log10 of *p*-values. **F-G)** Bioenergetic analysis of hBMSC using Seahorse analyzer with 6-8 technical replicates per group. F) The scatter plot shows the basal mitochondrial and glycolytic ATP production. The bar chart shows the mean percentage of mitochondrial and glycolytic ATP production. The box plot shows the total ATP production rate (mitochondrial and glycolytic). G) The scatter plot shows the basal mitochondrial respiration and glycolysis rates. The left box plot shows the change in the metabolic programme as the ratio between basal mitochondrial respiration and basal glycolysis. The middle box plot shows the basal glycolysis rates. The left panel show the total ATP production rate. **H)** Correlation between the percentage of heteroplasmy and mitochondrial function. Left panel: basal mitochondrial respiration rates. Center panel: basal glycolysis rates. Right panel: basal mitochondrial ATP production rates. Shadow shows a 95% confidence interval. n = 20 (10 m.3243A>G carriers and 10 controls matched by age, sex, and BMI). Box plot showed normalized data expressed as each individual’s mean of technical replicates. RNA sequencing and proteomics analysis were done in the same batch. Bioenergetic analysis of matched m.3243A<G carriers and controls hBMSC was done in the same Seahorse plate. The statistical analysis performed was Wilcoxon signed-ranked test. *: *p*-value < 0.05, **: *p*-value <0.01 ATP: adenosine triphosphate; BMI: body mass index; GO: Gene Ontology; hBMSC: human bone marrow mesenchymal stem cells; ddPCR: droplet digital polymerase chain reaction. OCR: oxygen consumption rate; ECAR: extracellular acidification rate

To study the effect of impaired translation of mDNA-encoded OXPHOS proteins in bone cells, we isolated hBMSCs from transiliac bone marrow aspirates. Carriers of pathogenic variant m.3243A>G have both mutated (m.3243A>G) and normal (wild-type) mDNA in each cell, defined as heteroplasmy. The variable level of heteroplasmy within cells and tissues correlates to some extent with the involvement of the respective organ in the disease development^19,20^. To evaluate heteroplasmy levels in hBMCS, we analysed the number of mDNA molecules carrying m.3243A>G and wild-type in hBMSCs by droplet digital PCR. Heteroplasmy ranged from 12-82% in m.3243A>G carriers, while the levels were below 0.1% in controls **(Figure 1C)**.

To explore the metabolic impacts of the pathogenic variant m.3243A>G in hBMSC, we performed a comprehensive analysis using RNA sequencing, discovery proteomics, and bioenergetic analysis that show an altered transcriptional, proteomic, and metabolic programming of carrierś hBMSCs **(Figure 1D-G)**. RNA sequencing of cultured hBMSCs showed 1,936 differentially expressed (DE) genes which were enriched mainly in mitochondrial functions, including mDNA translation, mitochondrial metabolism, morphology, and biogenesis **(Figure 1D)**. Unsupervised principal components analysis of the transcriptomic profile of hBMSCs separated carriers and controls into two groups with a small overlap between them **(Figure S1A)**. Furthermore, unsupervised cluster analysis of the 1,936 DE genes **(Figure S1B)** and 148 DE mitochondrial genes – according to MitoCarta 3.0^21^ – **(Figure S1C)** showed differences in the expression profile between the carriers and controls hBMSC groups. To further evaluate whether this transcriptional regulation is reflected in the protein levels, we performed large-scale discovery proteomics in hBMSC. The hBMSC proteome showed altered levels of metabolic proteins, specifically mitochondrial metabolism, such as electron transport chain (ETC), citric acid (TCA) cycle, and branched-chain amino acid catabolism (BCAA) **(Figure 1E)**. Overall, the transcriptome and proteome show altered regulation of mitochondrial metabolism in carrierś hBMSC compared to controls. One of the key roles of mitochondria is the production of cellular energy in the form of adenosine triphosphate (ATP), which is essential for highly active tissues such as bone. We found a highly glycolytic profile in carrierś hBMSCs, specifically increased glycolytic ATP production (FC = 1.48, p<0.01) and decreased mitochondrial ATP production (FC = 0.8) **(Figure 1F, Figure S2A-B, and Table S2)**. Notably, the net ATP production rate was unchanged, with a tendency towards a higher level in hBMSCs from carriers **(Figure 1F)**. This shift in bioenergetic phenotype highlights the increased glycolytic rates in carrierś hBMSC (FC = 1.56, p>0.01) **(Figure 1G)** in response to the decreased OXPHOS rate, as observed in the transcriptome, proteome and mitochondrial bioenergetics profiles **(Figure 1D-E, Figure S2A-J and Table S2).** Increased glycolytic bioenergetics has previously been observed in non-skeletal cells with compromised OXPHOS function, such as patients with mitochondrial disease^17,18,22^. The hBMSCs, like other stem cells, rely on glycolysis for self-renewal and proliferation^5^. This increased glycolytic profile in compromised OXPHOS cells could lead to stem cell exhaustion due to a long-term increased proliferation, with a decline in cell numbers, renewal capacity and bone formation capacity^23^.

Higher heteroplasmy levels of m.3243A>G is generally associated with an earlier debut of symptoms and a more severe clinical phenotype^19^. Accordingly, the levels of heteroplasmy correlated significantly with the metabolic program of carrierś hBMSC. The heteroplasmy level correlated negatively with metabolic programming index (mitochondrial respiration/glycolysis) (r = - 0.72, *p* = 0.02) and mitochondrial ATP production (r = - 0.84, *p* = 0.002) and positively with glycolytic ATP production (r = 0.84, *p* = 0.002) **(Figure 1H and Figure S3A-G)**. These findings align with previous results obtained from studies of cell cybrids showing that heteroplasmy percentage correlates with the upregulation of glycolytic genes^24^. The heteroplasmy percentage also correlated positively with the plasma lactate level **(Figure S3H)**. Furthermore, 98% of the 1,936 DE genes in carrierś hBMSC correlated with the percentage of heteroplasmy. On the contrary, heteroplasmy levels showed little correlation with the proteomic profiles (6.3%), indicating that posttranscriptional regulation is independent of the heteroplasmy level in carrierś hBMSCs.

Because the pathogenic variant m.3243A>G is located in the mDNA, we examined the effects of m.3243A>G in hBMSC on the expression of mDNA genes, which includes 37 genes: 2 rRNA, 22tRNAs, and 13 genes critical for the OXPHOS pathway. We detected the expression of 28 mDNA genes by RNA sequencing, with three significantly upregulated, one tRNA (*MT-TC*) and two OXPHOS (*MT-CO2* and *MT-CYB*) genes **(Figure S4A)**. At the same time, their protein levels were unaffected **(Figure S4)**. The *MT-TL1* gene, where m.3243A>G is located, was very low expressed in hBMSC **(Figure S4B)**. OXPHOS comprises more than 130 proteins mainly encoded in the nDNA, except for the 13 mDNA-encoded proteins^25^. The gene expression of 26% (34 genes) of nDNA-encoded OXPHOS proteins was upregulated; however, only two showed changes in the protein level **(Figure 2A)**. This upregulation of OXPHOS gene expression could reflect a compensatory mechanism to rescue the impaired mitochondrial ATP production in carrierś hBMSC **(Figure 1F-G)**. The increased OXPHOS gene expression is not reflected in increased protein levels, possibly due to low levels or misfolding of mDNA-encoded proteins caused by impaired mDNA translation in m.3243A>G carriers hBMSC. This negatively affects the ETC complexes’ stability, thus decreasing OXPHOS rates. OXPHOS and TCA activities are tightly coupled as oxidation of NADH and FADH2 by OXPHOS complexes I and II are required for the TCA cycle^13^. To evaluate the effects of the decreased OXPHOS activity observed in hBMSC from m.3243A>G carriers on the TCA cycle, we performed dynamic metabolic mapping using uniformly labelled glucose ([U-^13^C]glucose) to measure ^13^C-labeled cellular metabolites derived from glucose, which is the primary energy substrate source for bone^4,5^. The [U-^13^C]glucose is metabolised to [U-^13^C]pyruvate that can be converted into: **(1)** ^13^C-labelled lactate, **(2)** ^13^C-labelled alanine or **(3)** ^13^C-labelled citrate in the mitochondria via acetyl-CoA upon entry in the TCA cycle. hBMSC showed a low ^13^C enrichment in measured metabolites, indicating a slow glucose metabolism with a tendency to increased ^13^C-labeled lactate in m.3243A>G hBMSC compared to control. This is in line with the increased glycolytic rates observed in the bioenergetics assays **(Figure 2B and Figure 1F-G)**. Furthermore, we observed increased levels of rate-limiting glycolytic enzymes such as hexokinase 2 (HK2) and gamma enolase (ENO2), and lactate dehydrogenase isoforms (LDHA and LDHC) without changes in their gene expression, indicating a posttranslational regulation of glycolysis **(Figure 2C)**. Pyruvate enters the mitochondria and is metabolized to acetyl-CoA, which enters the TCA cycle. The relative abundance of intermediate TCA metabolites tended to be reduced in the first turn of the cycle (M+2 metabolites), while ^13^C-labeled malate and ^13^C-labeled fumarate were increased **(Figure 2B)**.

**Figure 2.**
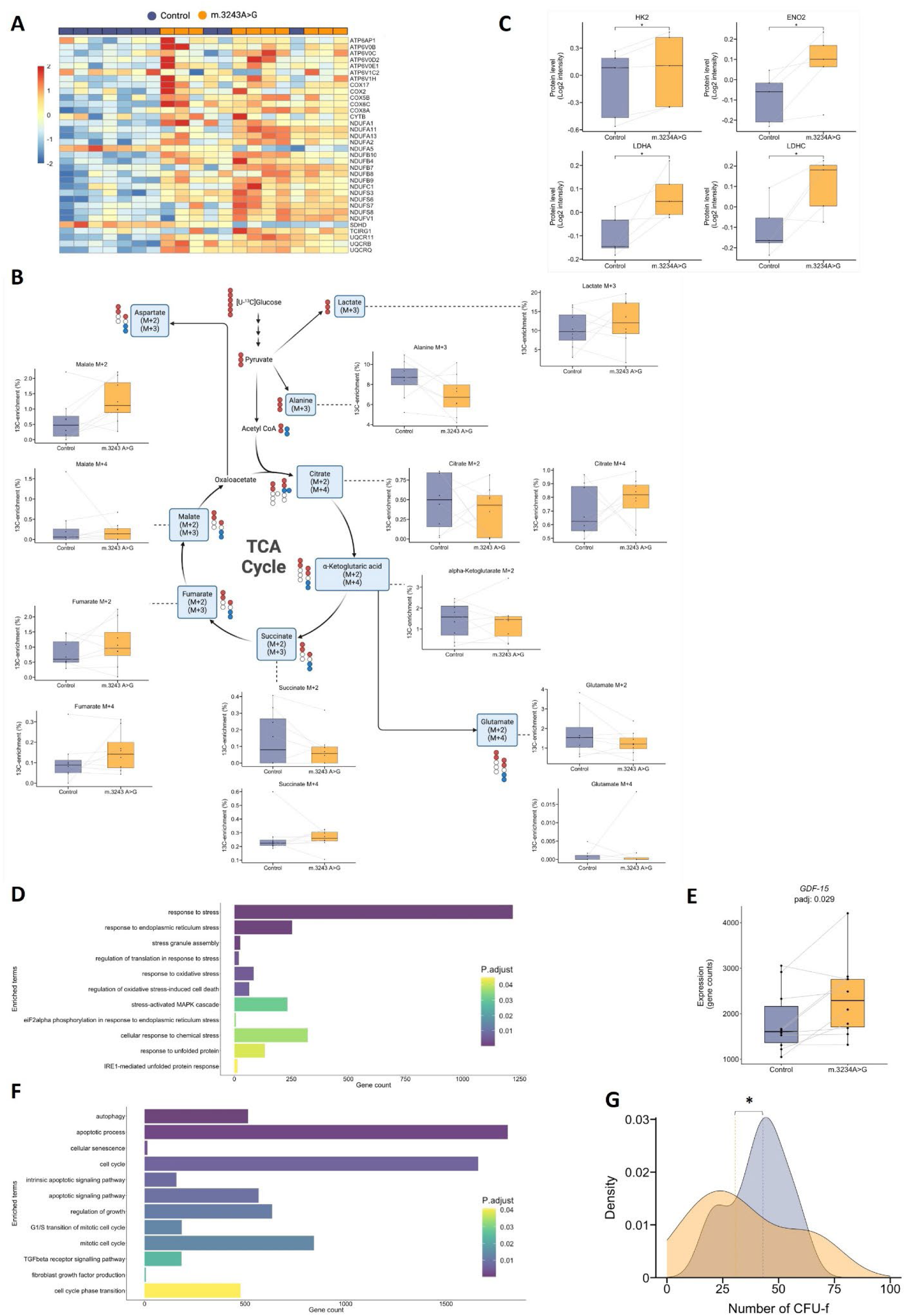
hBMSC from m.3243A>G subjects show a higher metabolic profile. **A)** Heatmap of the number of DE nDNA genes encoding OXPHOS proteins in hBMSC. **B)** Dynamic metabolic mapping of [U-^13^C]-glucose-derived metabolites in hBMSC after incubation with [U-^13^C]glucose analyzed by GC-MS. Alpha-ketoglutarate M+4 is not shown due to low detection levels. **C)** Protein levels of glycolysis-related proteins analysed by mass spectrometry-based proteomics. **D)** Enrichment analysis of DE genes related to MSR on hBMSC from m.3243A>G compared to controls using GO biological process terms as database. The enriched terms show the number of genes in each GO term and the adjusted *p*-value of the GO term. **E)** Boxplot of *GDF-15* gene expression levels (read counts) in hBMSC analysed by RNA sequencing. **F)** Enrichment analysis of DE genes related to autophagy, cell cycle, cellular proliferation, apoptosis and cellular senescence on hBMSC from m.3243A>G compared to controls using GO biological process terms as database. The enriched terms show the number of genes in each GO term and the adjusted *p*-value of the GO term. **G)** CFU-f formed from 0.5 x 10^6^ freshly isolated hBMSC during 14 days cultures, where m.3243A>G hBMSC showed decreased proliferation. n = 20 (10 m.3243A>G carriers and 10 controls matched by age, sex, and BMI). Box plot showed normalized data expressed as each individual’s mean of technical replicates. RNA sequencing, proteomics and metabolic mapping were done in the same batch. The statistical analysis performed was Wilcoxon signed-ranked test. *: *p*-value < 0.05 BMI: body mass index; CFU-f: colony forming units, fibroblasts; DE: Differentially expressed; ENO2: Gamma-enolase; ENO: GC-MS: gas-chromatography mass spectrometry; GDF-15: Growth Differentiation Factor 15-gene; GO: Gene Ontology hBMSC: human bone marrow mesenchymal stem cells; HK2: Hexokinase-2; LDHA: lactate dehydrogenase A; LDHC: lactate dehydrogenase C; MSR: mitochondria stress responses; OXPHOS: oxidative phosphorylation; TCA: tricarboxylic acid cycle; U-^13^C: 13 carbon glucose isotopomer

OXPHOS deficiency, even with normal ATP levels, triggers nuclear transcriptional responses such as unfolded protein response (UPR) and integrated stress response (ISR) to coordinately counteract the effects of OXPHOS dysfunction^22,26^. Here, we summarise these responses under the term mitochondrial stress response (MSR). MSR is dysregulated in hBMSC from m.3243A>G, as shown by the enriched terms related to stress responses in DE genes **(Figure 2D)**. Most of the genes involved in these pathways are upregulated, indicating an upregulation of the MSR in carrierś hBMSC. MSR activates the secretion of circulating cytokines such as growth/differentiation factor 15 (GDF-15) that impact whole-body energy metabolism^27^ and is considered a mitochondrial disease biomarker^28–30^. Carrierś hBMSC showed an increased expression of *GDF-15* (logFC = 0.36) **(Figure 2E)**. In addition, MSR activates a cellular transcriptional program, resulting in autophagy, cell cycle arrest, decreased proliferation, apoptosis or cellular senescence^31^. The transcriptional program of hBMSC from carriers showed enriched terms in these pathways **(Figure 2F)**. Overall, these transcriptional changes show activation of MSR in carrierś hBMSCs, which requires energetically demanding processes such as transcription, translation and secretion. These processes can act as an energy drain and, therefore, prevent other energy-costly processes such as cellular differentiation^14,22^. Cultured carrierś hBMSCs showed fewer colony-forming units-fibroblasts (CFU-f), indicating decreased proliferation and stemness **(Figure 2G)** that could be a sign of stem cell exhaustion due to the increased glycolytic profile **(Figure 1F-G).**

Collectively, this data shows that hBMSCs from m.3243A>G carriers present with impaired OXPHOS and an adaptative increased glycolytic profile, preserving total cellular ATP production. Impaired OXPHOS also activates the MSR transcriptional program in m.3243A>G hBMSC and the transcriptional program for autophagy, cell cycle arrest, decreased proliferation, apoptosis, and cellular senescence **(Figure 2D-H)**. Jointly, these changes could reduce the proliferation and stemness of the carriers’ hBMSCs, ultimately resulting in lower bone formation capacity.

### Highly glycolytic metabolic program in m.3243 A>G carrierś hBMSC decreases their osteogenic capacity

The importance of mitochondrial function for bone health has been shown in cellular and animal models, e.g. mitochondrial ATP production, TCA cycle intermediate metabolites, MSR activation,^5,32–34^ all of which are affected in hBMSCs from m.3243A>G carriers. Moreover, pharmacological inhibition of glycolysis promotes OXPHOS and increases BMD in mice^7^. We subsequently investigated if the highly glycolytic program of carrierś hBMSC decreased their ability to differentiate to osteoblasts and to form new bone. hBMSCs can differentiate into osteoblasts and adipocytes but are transcriptionally predisposed to osteoblast differentiation, which represses adipogenesis^35^. The hBMSC transcriptome of m.3243A>G carriers showed an altered gene expression of bone-related pathways; specifically, we observed a downregulation of transcription factors involved in bone formation, i.e. WWTR1 (also known as TAZ)^36,37^*, SMAD5*^38,39^*, SOX11*^40,41^ and *YAP1*^36,42^ **(Figure 3A-B)**. This led to an altered stem cell lineage determination in carrierś hBMSC compared to controls hBMSC **(Figure 3C-F)**. Carrierś hBMSC exhibited a decreased correlation between alkaline phosphatase (ALP) activity – an early osteogenesis marker – and age, while the controlś hBMSC showed no correlation **(Figure 3D)**. On the contrary, adipocyte differentiation capacity assessed by lipid droplet accumulation was not correlated with age in carrierś hBMSC but was correlated in controls **(Figure 3F)**. This difference in lineage determination pattern favourable to adipogenesis instead of osteogenesis has been observed in hBMSC from elderly population (> 85 years old), where an increase in adipogenesis is associated with decreased bone mass^43,44^. To further evaluate the hBMSC bone formation capacity *in vivo*, we performed a heterotopic assay with subcutaneous implantation of a mixture of hBMSC and hydroxyapatite – as a scaffold – in 8-weeks old immunodeficient female mice for 8 weeks. The hematoxylin-eosin staining of the implants revealed a decrease in the bone formation capacity of carrierś hBMSC compared to their matched controls **(Figure 3G)**.

**Figure 3.**
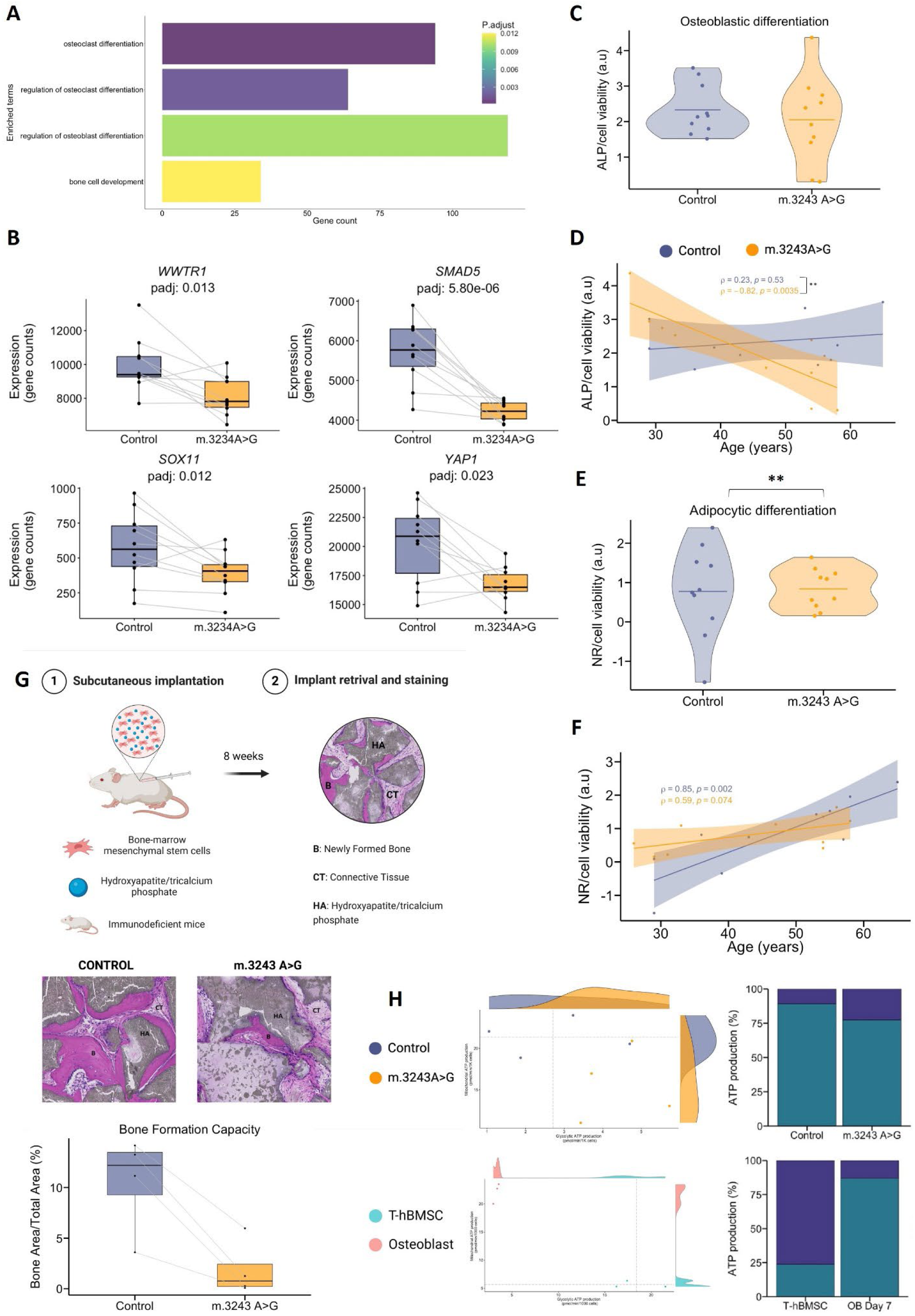
hBMSC from m.3243A>G show decreased bone formation capacity. **A)** Enrichment analysis of DE genes related to the skeleton on hBMSC from m.3243A>G compared to controls using GO biological process terms as database. The enriched terms show the number of genes in each GO term and the adjusted *p*-value of the GO term. **B)** Box plot with gene expression levels (read counts) of transcription factors that promote bone formation analysed by RNA sequencing. **C)** Early osteogenic marker, ALP activity was analyzed by colourimetry and normalized to cell viability to account for differences in cell proliferation. hBMSC were differentiated to OB for 7 days in osteoblast induction media. **D)** Correlation between age and ALP activity normalized to cell viability. Shadow shows a 95% confidence interval. **E)** Lipid droplets, adipogenic marker, were analysed by Nile Red fluorescent dye using high image content analysis, Operetta. Nile Red data is shown as MFI (Mean Fluorescent Intensity) normalised to cell viability to account for differences in cell proliferation. The *p*-value reflects the Bartlett Test of Homogeneity of Variances. **F)** Correlation between age and Nile Red (MFI/cell viability). Shadow shows 95% confidence interval. **G)** Heterotropic bone formation assay using hBMSC and hydroxyapatite/tricalcium phosphate as a scaffold, implanted subcutaneously in immunodeficient mice for 8 weeks. Left panel: representative histology section stained with HAE. Newly formed bone is purple. Connective tissue is pink. Hydroxyapatite/tricalcium phosphate is grey. Right panel: Quantification of the newly formed bone by analysing bone area (purple) to total area (purple + pink). **H)** Bioenergetic analysis of osteoblasts using Seahorse analyzer with 6-8 technical replicates per group. The scatter plots show the basal mitochondrial and glycolytic ATP production. The upper scatter plot shows the osteoblasts differentiated from carrierś hBMSC and matched controls (n = 8, 4 carriers and 4 controls matched by age, sex and BMI). The lower scatter plot shows the undifferentiated T-hBMSC and osteoblasts differentiated from T-hBMSC for 7 days (n = 3 independent experiments). n = 20 (10 carriers and 10 controls matched by age, sex, and BMI). Box plot showed normalised data expressed as each individual’s mean of technical replicates. RNA sequencing and heterotropic bone formation were done in the same batch. The statistical analysis performed was Wilcoxon signed-ranked test. *: *p*-value < 0.05. **: *p*-value<0.01. ALP: alkaline phosphatase; B: trabecular bone; CFU-f: colony forming units, fibroblasts; CT: connective tissue; HA: hydroxyapatite/tricalcium phosphate; HAE: hematoxylin-eosin; hBMSC: human bone marrow; OB: osteoblast; SMAD5: SMAD family member 5-gene; SOX11: SRY-BOX 11-gene; T-hBMSC: telomerized hBMSC; WWTR1: WW domain-containing transcription regulator 1-gene; YAP1: YES- associated transcriptional regulator.

These data show that hBMSC from m.3243A>G carriers have decreased *in vivo* bone formation capacity despite unchanged total cellular energy production. This suggests that m.3243A>G hBMSC cannot modulate OXPHOS activity to meet the demands for bone formation, despite the compensatory upregulation of glycolysis. This high dependency on a glycolytic bioenergetic program to maintain energy production levels probably leads to impaired metabolic flexibility overall impairing bone formation. Metabolic flexibility is defined as the capacity of the cells to adapt to changes in energy demands, e.g. increasing mitochondrial ATP production^45^. Whether the increased energy demand for osteoblastś differentiation is fulfilled by increasing mitochondrial or glycolytic ATP production remains controversial. Studies on cultured human cells mainly show increased mitochondrial ATP production, while rodent studies primarily reveal increased glycolytic ATP production^5^. To provide further evidence of mitochondrial ATP production being key to human cells, we analysed the bioenergetic profile of a subset of carrierś hBMSC after differentiation to osteoblasts for 12 days. We observed a shift to an increased OXPHOS bioenergetic profile in osteoblasts from both carriers and controls, with carriers’ osteoblasts showing a smaller increase in mitochondrial ATP production (FC = 0.72) and remaining more glycolytic (FC = 1.62) than controlś osteoblasts **(Figure 3H)**. This suggests that impaired metabolic flexibility in carrierś hBMSC caused by a high dependence on a glycolytic bioenergetic profile impairs their ability to differentiate into mature osteoblasts.

To evaluate whether the increased glycolytic rate or the impaired OXPHOS limits the hBMSC bone formation capacity, we analysed the bioenergetic profile of a telomerized hBMSC (T- hBMSC), a cell line with high proliferation rates and high bone formation capacity developed in our laboratory^46^. T-hBMSCs have a higher glycolytic profile than carrierś hBMSC; however, upon osteoblast differentiation, T-hBMSCs decrease glycolysis within 24 hours of osteoblastogenesis induction while maintaining low mitochondrial respiration. Subsequently, T-hBMSC progressively increased their OXPHOS rates and mitochondrial ATP production, peaking at day 7 **(Figure 3H)**. Thus, the decreased osteogenic capacity of carrierś hBMSC is highly likely caused by their impaired OXPHOS capacity.

### Osteoclasts from m.3243A>G show increased TRAcP activity without changes in mitochondrial function

Bone remodelling is a dynamic and coordinated process between bone formation by osteoblasts and bone resorption by osteoclasts. Osteoclasts are derived from the hematopoietic progenitors in bone marrow and have a high abundance of mitochondria^47^. While osteoclast differentiation depends on OXPHOS, glycolysis is the main bioenergetic pathway for bone resorption activity *in vitro*^47^. To further evaluate the effects of the m.3243A>G impaired OXPHOS on bone resorption, we studied osteoclasts derived from peripheral blood mononuclear cells (PBMCs) from m.3243A>G carriers and their matched controls **(Figure 4A)**. We and others have previously reported that the variant m.3243A>G is selected against with every cellular division; thus, replicative cells like monocytes show decreased heteroplasmy level with age^48,49^. Accordingly, matured osteoclasts (day 9 of differentiation) from carriers generally showed lower heteroplasmy levels (7 - 44%) than hBMSCs **(Figure 4B)**. The heteroplasmy of osteoclasts correlated positively with the heteroplasmy of hBMSCs (R = 0.82) **(Figure 4C)**. Although the bioenergetic profiles of carrierś mature osteoclasts were not different than those of their matched controlś mature osteoclasts **(Figure 4D-F and Table S3)**, matured osteoclasts reseeded on bone slices showed increased bone resorption activity for six of the ten carrierś mature osteoclasts **(Figure 4G-H and Figure S5)**. This indicates that other factors besides bioenergetics influences bone resorption in carrierś mature osteoclasts. At the same time, we observed altered resorption patterns with respect to the correlation between the percentage of pits (round cavities made by immobile osteoclasts) and trenches (elongated resorption made by osteoclasts moving across the surface) surface per eroded surface (PS/ES and TS/ES)^50^. Mature osteoclasts from carriers showed no correlation between pits and trenches surfaces per eroded surface as was observed with the mature osteoclasts from the controls **(Figure 4I)**. This suggests a decreased mobility of carrierś osteoclasts during bone resorption. Pit-forming osteoclasts are associated with increased eroded surface compared to trenches-forming osteoclasts^50^. During bone resorption, osteoclasts secrete the tartrate-resistant acid phosphatase (TRAcP) enzyme to aid in the degradation of bone tissue^51^. The activity of secreted TRAcP during bone resorption was significantly higher in m.3243A>G carriers (FC 1.41) than in controls **(Figure 4J)**. Moreover, the activity of secreted TRAcP during differentiation (from day 2 to day 9) was also significantly increased in osteoclasts from m.3243A>G carriers **(Figure 4K)**. This, together with a tendency to an increased number of nuclei per osteoclast **(Figure 4L)**, indicates a higher differentiation process of carrierś osteoclasts. None of these parameters correlated with the heteroplasmy levels or the bioenergetic profile (data not shown), indicating that other parameters drive these changes. Studies on osteoclasts with mDNA pathogenic variants caused by deficient mDNA replication in the PolgA^mut/mut^ mice – so-called mutator mice – showed increased osteoclast resorptive activity and bone loss^52^. The mutator mice lack mDNA proofreading activity^53^ leading to severe OXPHOS dysfunction and increased osteoclast activity^52^. In the present study, osteoclasts from m.3243A>G carry a single pathogenic nucleotide variant with low levels of heteroplasmy, which could explain the normal OXPHOS activity and why there is only a slight increase in resorptive and secreted TRAcP activity.

**Figure 4.**
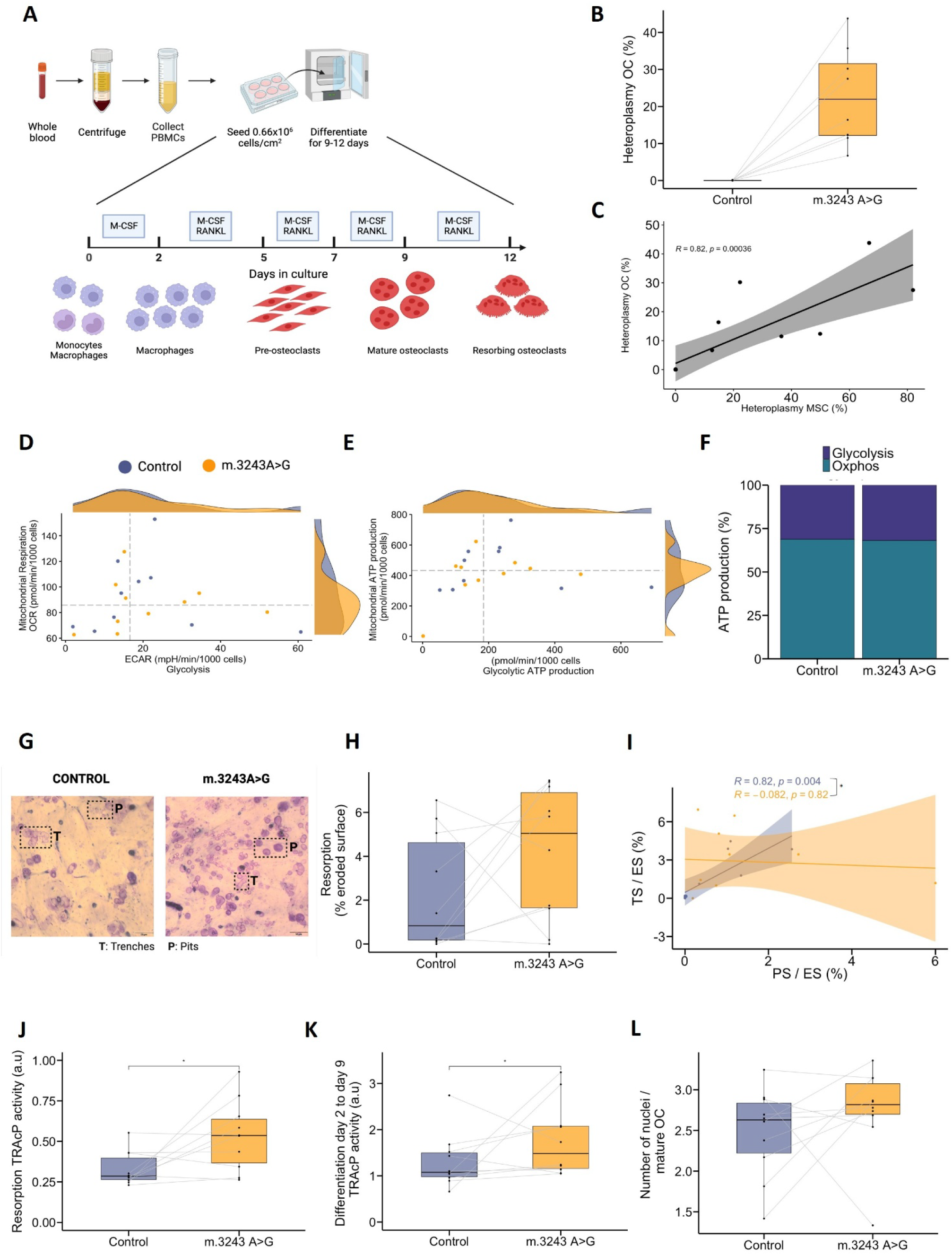
m.3243 A>G osteoclasts show normal mitochondrial function. **A)** Schematic representation of the OCs experiments. Peripheral blood mononuclear cells were isolated from peripheral blood and cultures with 25 ng/mL M-CFS until day 2, when 25 ng/mL RANKL was added. Cell culture media was collected at each media change for TRAcP activity. OCs were considered mature at day 9 when mitochondrial function analysis using Seahorse Extracellular analyzer was done, DNA was collected, and OCs were re-seeded onto bone slices for resorptive activity. **B)** Heteroplasmy analysis of mature OCs was done using digital PCR. **C)** Pearson correlation of heteroplasmy in mature OCs and hBMSCs from m.3243A>G carriers. Grey shadow shows 95% confidence interval. **D-F)** Mitochondrial function analysis of mature OCs using Seahorse analyser (6-8 wells/group). The scatter plot shows the basal mitochondrial respiration and glycolysis of mature OC **(D)** and basal mitochondrial and glycolytic ATP production **(E)**. The bar chart shows the proportion of mitochondrial ATP and glycolytic ATP **(F)**. **G)**Representative pictures of bone slices stained with toluidin blue show the resorption activity (pits and trenches) of mature OCs after 3 days of culture. Examples of pits and trenches are marked. **H)** Resorption activity of mature OCs quantified as the percentage of eroded surface by bone Surface (eight replicates). **I)** Pearson correlation between the percentage of TS/BS and the percentage of PS/BS. The shadow shows 95% confidence interval. **J)** Extracellular TRAcP activity of mature OCs on bone slices (eight replicates). **K)** Extracellular TRAcP activity of cells under RANKL stimulation for differentiation analysed by colorimetry. **L)** Number of nuclei per mature OCs quantified by brightfield imaging from 12 images across a 75-cm^2^ culture flask. n = 20 (10 m.3243A<G carriers and 10 controls matched by age, sex, and BMI). Except for heteroplasmy analysis, where n = 16 (8 m.3243A<G carriers and 8 controls matched by age, sex, and BMI). Box plot showed data expressed as each individual’s mean of technical replicates. The statistical analysis performed was Wilcoxon signed-ranked test. * p-value < 0.05 ATP: adenosine triphosphate; ECAR: extracellular acidification rate; M-CFS: Macrophage colony-stimulating factor; RANKL: Receptor activator of nuclear factor kappa-Β ligand; OC: osteoclast; OCR: oxygen consumption rate; PS/BS: pits surface/bone Surface; TRAcP: Tartrate-resistant acid phosphatase; TS/BS: trenches surface/bone surface

### Bone biopsies from m.3243A>G carriers indicate a delay in bone formation

The *in vivo* clinical effects of the mitochondrial dysfunction on bone tissue were also evaluated in transiliac bone biopsies from seven m.3243A>G carriers and an equal number of healthy individuals matched on age and sex, but not BMI that was higher in the controls than in the carriers. Micro-computed tomography (µCT) of the bone biopsies showed a 50% lower trabecular bone volume (BV/TV) that was associated with a lower trabecular thickness **(Figure 5A-F and Figure S6)**. No difference was observed in the cortical bone parameters between carriers and matched controls **(Figure 5G and Figure S6)**. Bone histomorphometry of the trabecular bone surfaces showed no difference between carriers and controls in eroded surface (ES) per bone surface (ES/BS) – reflecting the initial resorption and reversal-resorption phase bone –, as well as osteoid surface (OS) per bone surface (OS/BS), and mineralising surface (MS) per bone surface (MS/BS) – both reflecting the bone formation phase **(Figure 5H-J and M)**. However, the extent of ES without neighbouring OS was increased in carriers, indicating a slight delay in the initiation of bone formation on ES **(Figure 5K-L)**. Furthermore, the mineral apposition rate (MAR) could only be measured in four of the m.3243A>G carriers, as the double tetracycline labelling was missing in the remaining carriers, indicating a decreased MAR that reflects a reduced rate of mineralisation once the process has started **(Figure 5M-N)**. On the contrary, adipocyte histomorphometry showed a tendency to increased mean adipocyte diameter (Ad.Pf.Dm), with unchanged adipocyte area (Ad.Ar/Ma.Ar) and density (Ad.Pf.N/Ma.Ar) **(Figure 5O and Figure S7)**.

**Figure 5.**
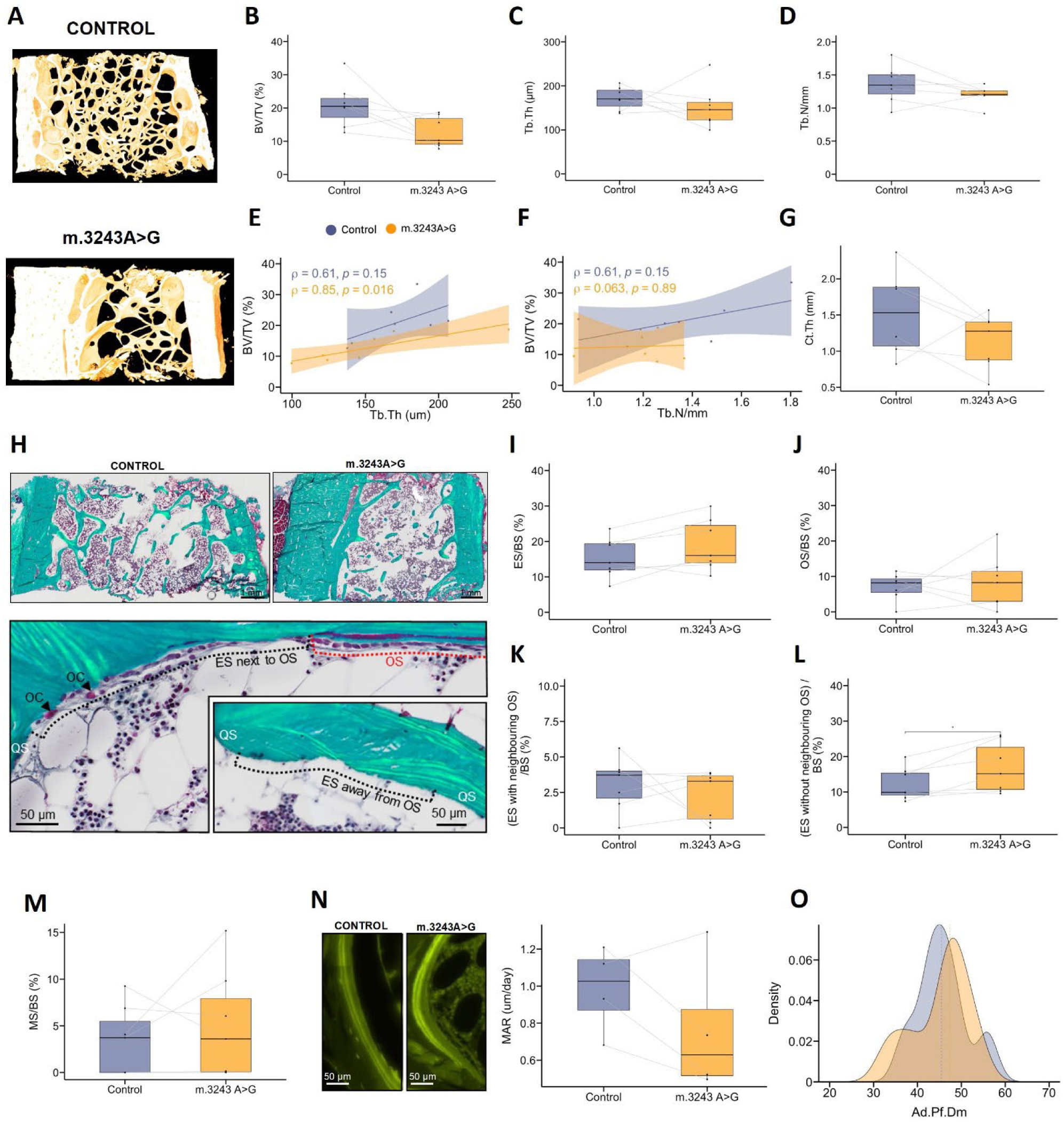
Delayed bone formation in iliac crest bone biopsies from m.3243 A>G carriers. **A)** Representative reconstructed µCT images of transiliac bone biopsies. **B-D)** Trabecular bone microarchitecture determined by µCT. Bone density determined as BV/TV (bone volume / total volume) **(B)**, Tb.Th = trabecular thickness **(C)**, Tb.N = trabecular number **(D)**. **E-F)** Spearman correlation analysis between BV/TV (%) and trabecular parameters: Tb.Th **(E)** and Tb.N **(F)**.The shadow shows 95% confidence interval. Cortical bone microarchitecture determined by µCT. Representative image of the histology analysis. **I-J)** Extension of the eroded surface (ES) **(I)** and the osteoid surface (OS) **(J)** related to the total bone surface (BS). **K-L)** The distance between the ES and the OS. Mineralization Surface related to total BS. Tetracycline labelling to mark day 0 and day 12 of bone formation by green fluorescence. The distance between both green fluorescent lines indicates the amount of bone formation in 12 days. Right panel shows quantification of bone formation analysed as Mineral Apposition Rare (MAR). Mean adipocyte profile diameter (Ad.Pf.Dm). n = 14 (7 carriers and 7 controls matched by age and sex), except MAR where n = 8 (4 carriers and 4 controls matched by age and sex). The statistical analysis performed was Wilcoxon signed-ranked test. * *p*-value < 0.05. Ad.Dm: adipocyte diameter; BS: bone surface; BV: bone volume; Ct.Th: cortical thickness; ES: eroded surface; MAR: mineral apposition rate; OS: osteoid surface; Tb.Th: trabecular thickness; Tb.N: trabecular number; TV: total volumen.

To further evaluate the effect of these microarchitecture changes on the macro-architecture of the bone, we performed DXA scans on all 20 study participants. BMD values were numerically but not significantly lower at the lumbar spine, femoral neck, and total hip **(Table S4)**. We previously reported lower BMD in a substantially larger m.3243A>G cohort^16^, which aligns with numerically lower BMD in m.3243A>G carriers in the present study. Considering that mechanical loading due to body weight is a key regulator of bone mass, we cannot exclude the possibility that the differences in BV/TV observed reflect the lower BMI in m.3243A>G carriers compared to controls rather than impaired bone formation.

Overall, these results indicate that the decreased bone formation capacity observed in cultured hBMSC from m.3243A>G carriers could be reflected in the bone tissue of these individuals, who have lower BMD in the trabecular bone, which is more metabolically active than cortical bone.

### Treatment with KH183 promoted a bioenergetically more efficient OXPHOS in m.3243A>G carrierś hBMSC

Treatments for mitochondrial diseases are in continuous development. Currently, a new treatment, Sonlicromanol, targeted for patients carrying the pathogenic variant m.3243A>G is nearing phase III clinical trial^54^. Sonlicromanol is a redox modulator with anti-inflammatory properties that counteracts oxidative stress, lipid peroxidation and ferroptosis, a common consequence of impaired OXPHOS function^55^. To evaluate whether rescuing OXPHOS impairment in hBMSC from m.3243A>G carriers improves bone formation, we treated hBMSC with KH183 – a pharmacologically active metabolite of Sonlicromanol. After three days of treatment with KH183, we observed a restoration of the transcriptional, proteomic, and metabolic program of m.3243A>G carrierś hBMSCs with increased mitochondrial respiration and decreased glycolysis **(Figure 6)**. RNA sequencing analysis revealed 6,846 DE genes after KH183 treatment that were enriched mainly in metabolic pathways and mitochondrial functions. Enriched terms in mitochondrial functions showed an overall downregulation, including morphological changes, mitochondrial ATP production, mitochondrial-induced apoptosis and mitophagy **(Figure 6A and S9A)**. This downregulation of the mitochondrial transcriptome indicates an improvement in mitochondrial functions since untreated carrierś hBMSC showed an upregulated mitochondrial transcriptome as a potential compensatory mechanism for OXPHOS deficiency **(Figure 1D)**. Furthermore, we observed transcriptional upregulation of 7 mDNA genes encoding proteins from complex I and IV and the two mDNA-encoded ribosomal RNAs (rRNA) while their protein levels were unaffected **(Figure S8)**. On the contrary, 38% (49 genes) of OXPHOS genes encoded in the nDNA showed downregulation, while only 9 OXPHOS proteins showed changes in the protein level **(Figure S9B)**. This transcriptional program was associated with a metabolic switch of increased mitochondrial ATP production and decreased glycolytic ATP production, supported by decreased gene expression of glycolytic pathway **(Figure 6B-D and Figure S10)**. However, total ATP production remained unchanged **(Figure 6E)**. Treatment with KH183 restored the bioenergetic profile of the carrierś hBMSC – 50% mitochondrial and 50% glycolytic ATP production with a negative correlation between them **(Figure 6D-F)**. The restored bioenergetic profile is similar to the bioenergetic profile of the controls hBMSC **(Figure 1F-G)**. The bioenergetically more efficient OXPHOS metabolic profile induced by KH183 could be caused by the altered composition of the OXPHOS complexes promoting OXPHOS rather than glycolysis. A key regulator of OXHPOS functional organization is the mitochondrial cytochrome oxidase 7 (COX7A) isoforms. COX7A isoforms promote the functional reorganization of two distinct mitochondria respiratory chain (MRC) structures: C-MRC, which is more efficient OXPHOS bioenergetics, and S-MRC, which is more glycolytic bioenergetics by increasing COX7A1 or COX7A2, respectively^56^. The gene expression of COX7A isoforms changes upon KH183 treatment by increasing COX7A1 and decreasing COX7A2, indicating an abundance of C-MRC structures promoting a more OXPHOS metabolism in m.3243A>G carrierś hBMSC **(Figure 6G)**. Furthermore, the third COX7A isoform, COX7A2L – known as “SC-associated factor 1” (SCAFI) – promotes a more glycolytic metabolic program^56^. In line with the metabolic reprogramming of KH183 treatment, COX7A2L gene expression is decreased upon KH183 treatment in carrierś hBMSC **(Figure 6G)**. The COX7A-isoform-dependent MRC organisation is tightly regulated by the pyruvate dehydrogenase complex (PDH) activity that provides acetyl-CoA to enter the TCA cycle, providing substrates for OXPHOS. PDH activity is regulated by phosphorylation inhibition by pyruvate dehydrogenase kinases (PDKs)^56^. The most expressed *PDK* isoforms in the bone marrow are *PDK1* and *PDK3*, that are downregulated after treatment with KH183, thus promoting PDH activity **(Figure 6H)**. Overall, this indicates that KH183 affects changes in the bioenergetics transcriptional program of carrierś hBMSC, leading to an increased abundance of C-MRC structures that are more OXPHOS efficient and, thus, restoring their metabolic program. The percentage of heteroplasmy correlates with 99% (6794 out of 6846) of the DE genes in hBMSC. In comparison, heteroplasmy levels showed little correlation with the proteomic profiles (6.9%) of carrierśhBMSC. This indicates that KH183 regulates the transcriptome of carrierś hBMSC according to the heteroplasmy levels, while their proteome regulation is independent of heteroplasmy.

**Figure 6.**
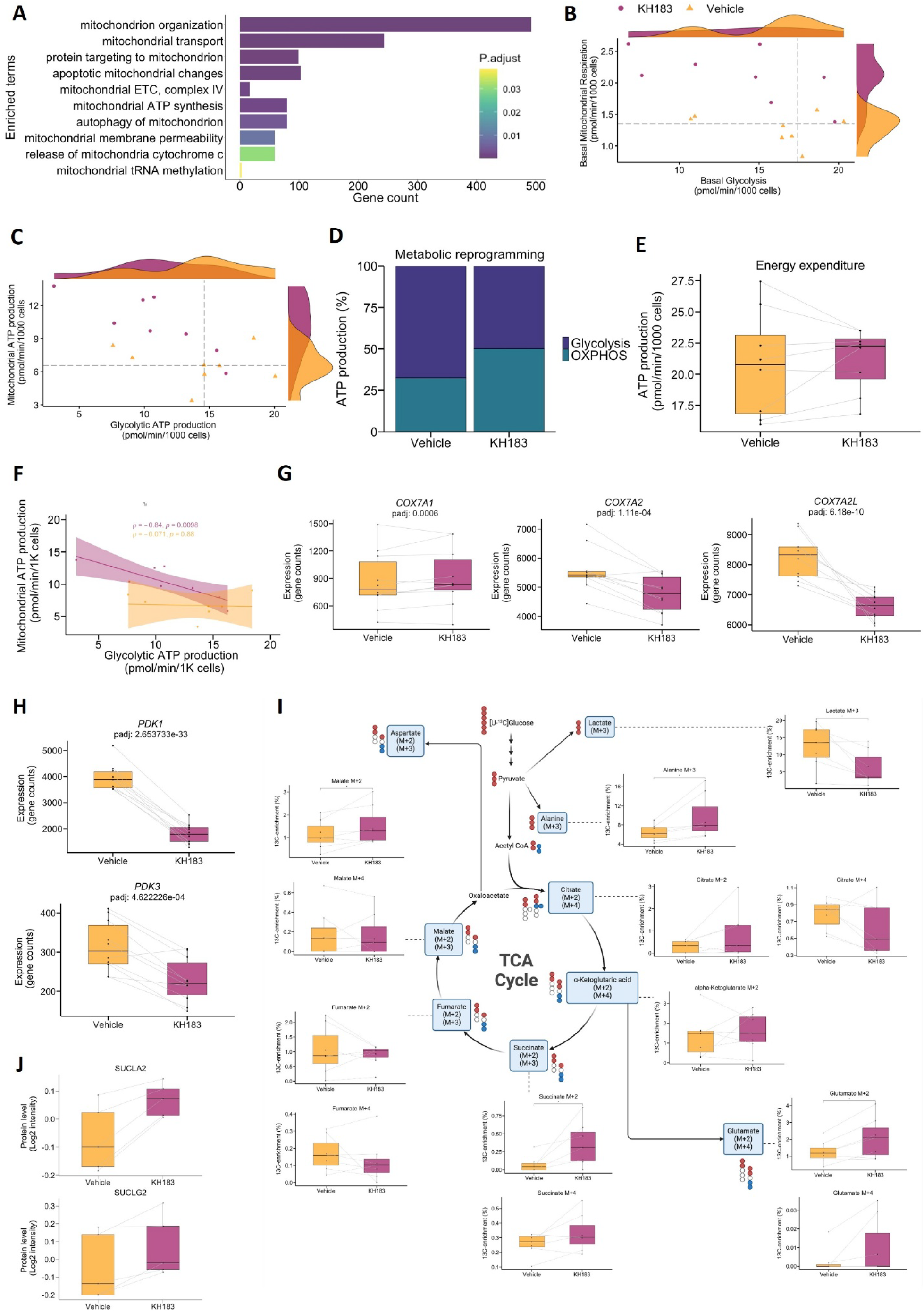
KH183 restores the metabolic program of hBMSC from m.3243A>G. hBMSC were cultured for 3 days with KH183 (5 uM) or DMSO as vehicle. **A)** Enrichment analysis of DE genes related to mitochondrial function in hBMSC from m.3243A>G compared to controls using GO biological process terms as database. The enriched terms show the number of genes in each GO term and the adjusted *p*-value of the GO term. **B-F)** Mitochondrial function analysis of hBMSC using Seahorse analyzer. The scatter plot shows the basal mitochondrial respiration and glycolysis **(B)** and basal mitochondrial and glycolytic ATP production **(C)**. The proportion of mitochondrial ATP and glycolytic ATP production **(D)**. Total ATP production **(E)**. Spearman correlation of mitochondrial ATP and glycolytic ATP production **(F)**. **G)** Box plot with gene expression levels (read counts) of *COX7A* isoforms analysed by RNA sequencing. **H)**Box plot with gene expression levels (read counts) of *PDK* isoforms analysed by RNA sequencing. **I)** Dynamic metabolic mapping of [U-^13^C] glucose-derived metabolites in hBMSC analysed by GC-MS. Alpha-ketoglutarate M+4 is not shown due to low detection levels. **J)** Protein levels of TCA cycle-related proteins measured by mass spectrometry. n = 16 (bioenergetics/RNAseq) and n =14 (metabolic mapping) m.3243A>G carrierś hBMSC treated with KH183 or DMSO. Box plot show normalised data expressed as each individual’s mean of technical replicates. RNA sequencing, proteomics and metabolic mapping were done in the same batch. Bioenergetic analysis of matched m.3243A<G carriers and controls hBMSC was done in the same Seahorse plate. The statistical analysis performed was Wilcoxon signed-ranked test. *: p-value < 0.05 *COX7A*: COX7A2-like gene; DMSO: Dimethyl sulfoxide; GC-MS: gas-chromatography mass spectrometry; GO: Gene Ontology; hBMSC: human bone marrow mesenchymal stem cell; OXPHOS: oxidative phosphorylation; *PDK*: pyruvate dehydrogenase kinase; TCA: tricaboxylic acid

Oxidation of nutrients – pyruvate, fatty acids, and amino acids – by OXPHOS is key for bone formation not only for ATP production, but also to produce acetyl-CoA and TCA cycle metabolites^5,57^. To evaluate whether KH183-induced increased OXPHOS activity affects the TCA cycle and overall glucose metabolism, we incubated the cells with [U-^13^C]glucose and analysed ^13^C-labelled metabolites. After KH183 treatment, the levels of ^13^C-labelled lactate significantly decreased in line with the decreased glycolytic rates observed, while ^13^C-labelled alanine was increased **(Figure 6I)**. ^13^C-labelled succinate was significantly increased after treatment, and the other ^13^C-labelled TCA metabolites tended to increase levels, indicating an increase in first-turn TCA cycle intermediates **(Figure 6I)**. Furthermore, SUCLA2 and SUCLG2 protein – involved in the production of succinate – levels were also increased after KH183 treatment, overall indicating a regulation of the succinate, a TCA cycle metabolite that directly connects it with OXPHOS via the complex II (succinate dehydrogenase, SDH) which converts succinate into fumarate^13^.

Overall, these data indicate a restoration of the energy metabolic profile of carrierś hBMSC after KH183 treatment by restoration of the mitochondrial transcriptome to the level of the controls. This change of the mitochondrial transcriptome improves mitochondrial ATP production and decreases the mitochondrial responses to OXPHOS deficiency. After treatment with KH183, we observed an overall downregulation of the mitochondrial pathways in carrierś hBMSC that were upregulated in untreated hBMSC **(Figure 1D and Figure 6A),** leading to a general downregulation of MSR genes **(Figure 7A)** together with a downregulation of mitochondrial disease biomarker *GDF-15* (LogFC = - 0.58) **(Figure 7C).** Downregulation of MSR led to a downregulation of DE genes involved in autophagy, cell cycle arrest, decreased proliferation, apoptosis, and cellular senescence **(Figure 7B)**. This improved mitochondrial function associated with the downregulation of genes involved in cellular stress responses, indicates a healthier cellular status in carrierś hBMSC.

**Figure 7.**
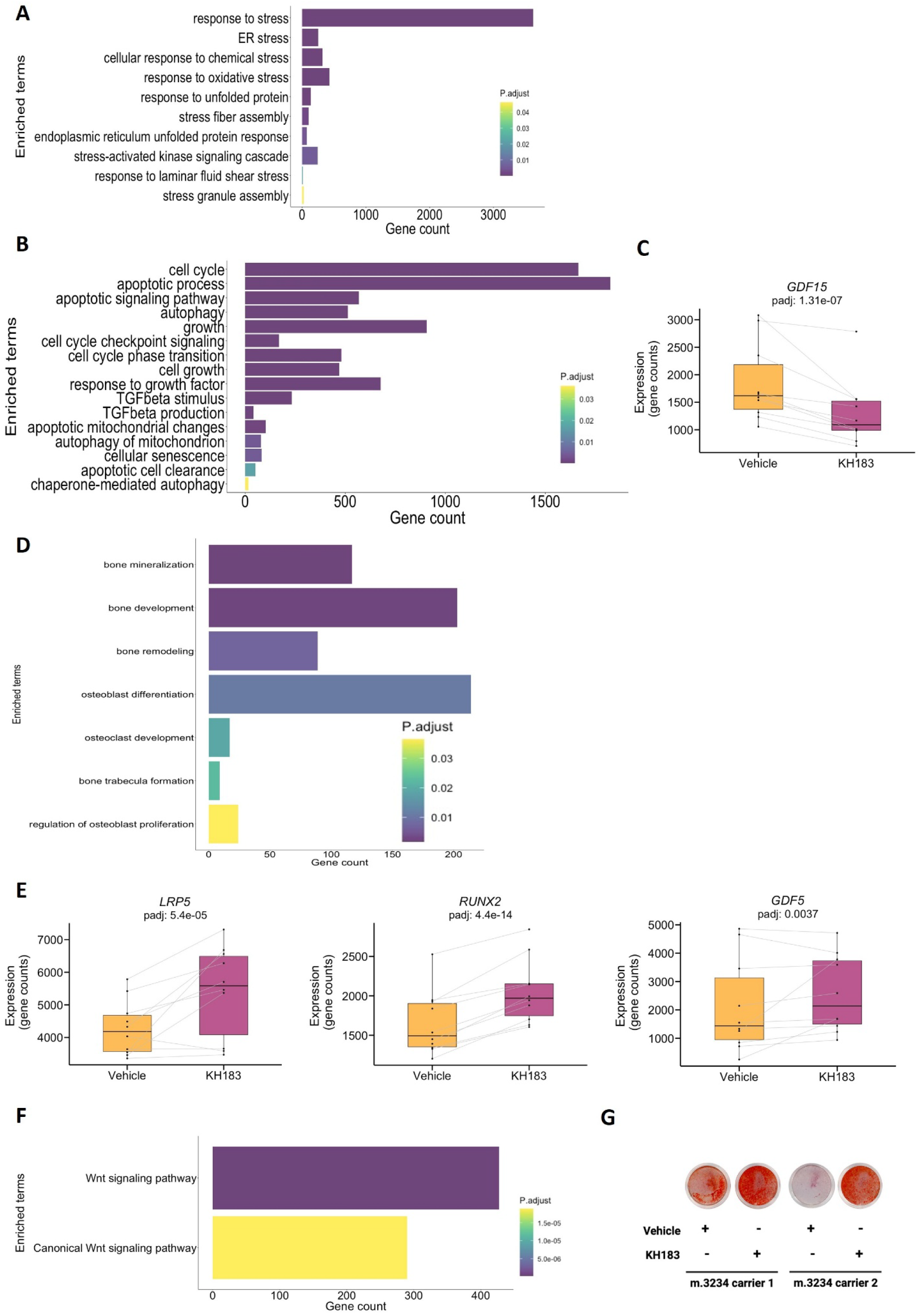
Improved mitochondrial function by KH183 alters the transcriptional program of hBMSC from m.3243A>G carriers. hBMSC were cultured for 3 days with KH183 (5 uM) or with DMSO (Vehicle). **A-B**) Enrichment analysis of DE genes related to MSR (A) and to autophagy, cell cycle, cellular proliferation, apoptosis and cellular senescence (B) in hBMSC from m.3243A>G after KH183 treatment compared to vehicle (DMSO) using GO biological process terms as database. The enriched terms show the number of genes in each GO term and the adjusted *p*-value of the GO term. **C)** Boxplot of *GDF-15* gene expression levels (read counts) in hBMSC analysed by RNA sequencing. **D)** Enrichment analysis of DE genes related to osteogenesis in hBMSC from m.3243A>G after KH183 treatment compared to vehicle (DMSO) using GO biological process terms as database. The enriched terms show the number of genes in each GO term and the adjusted *p*-value of the GO term. **E)** Boxplot of osteogenic gene expression levels (read counts) in hBMSC analysed by RNA sequencing. **F)** Enrichment analysis of DE genes related to Wnt pathway in hBMSC from m.3243A>G after KH183 treatment compared to vehicle (DMSO) using GO biological process terms as database. The enriched terms show the number of genes in each GO term and the adjusted *p*-value of the GO term. **G)** Mineralization capacity of hBMSC stained with Alizarin Red after 12 days in osteoblast induction media and KH183/vehicle (DMSO) treatment. n = 16 (bioenergetics/RNAseq) and n =14 (metabolic mapping) m.3243A>G carrierś hBMSC treated with KH183 or vehicle (DMSO). Box plot showed normalized data expressed as each individual’s mean of technical replicates. RNA sequencing, proteomics and metabolic mapping were done in the same batch. The statistical analysis performed was Wilcoxon signed-ranked test. *: p-value < 0.05 DMSO: Dimethyl sulfoxide; GDF5: growth differentiation factor 5; hBMSC: human bone marrow mesenchymal stem cell; LRP5: low density lipoprotein receptor-related protein 5; RUNX2: Runt-related transcription factor 2.

### More efficient OXPHOS in m.3243A>G carrierś hBMSC associates with a bone-forming transcriptional program

Previous studies have shown that several mitochondrial functions are important for osteoblastogenesis^5,32–34^. KH183 treatment improved key aspects of mitochondrial function in in carrierś hBMSC including OXPHOS activity, MSR, and mitophagy **(Figure 6 and Figure 7A- B)**. Thus, we hypothesized that treatment with KH183 can improve the osteogenic capacity of carrierś hBMSC. In line with our hypothesis, we observed upregulation of DE genes associated with bone formation **(Figure 7D)**. Moreover, key genes important to osteoblastogenesis such as *LRP5*, *RUNX2*, *TGFB3*, and *GDF5*, were upregulated in carrierś hBMSC after KH183 treatment **(Figure 7E)**. LRP5 is a Wnt receptor that enables Wnt activation, a key signalling pathway for osteoblast differentiation and bone formation^58,59^. Upregulation of *LRP5* gene expression was associated with an upregulation of Wnt pathway genes in KH183-treated carrierś hBMSC **(Figure 7F)**. Several studies show the Wnt-mitochondria axis, where inhibition of Wnt signalling impairs mitochondrial ATP production and viceversa^60,61^. Extending these findings, we show that restoring mitochondrial ATP production increases the expression of Wnt-associated genes in OXPHOS-deficient hBMSC. Importantly, we observed that KH183 improved the mineralisation capacity of m.3243A>G hBMSCs **(Figure 7G**), further supporting that promoting OXPHOS in m.3243A>G hBMSCs increases bone formation. While pharmacological inhibition of glycolysis and a subsequent increase in OXPHOS resulted in higher mineral density in mice^7^, the present study demonstrates that OXPHOS can be targeted for enhancing bone formation in humans.

## Discussion

Bone remodelling is essential for developing and maintaining bone mass, structure, and strength. Considering that this process is highly energy demanding, we investigated whether inherited impaired OXPHOS – causing decreased mitochondrial ATP production – would impede bone cell function and key aspects of bone health in adult men and women. While ATP requirements were compensated by increased glycolytic activity in hBMSCs from individuals with impaired OXPHOS (m.3243A>G carriers) **(Figure 1F-G)**, this change in the bioenergetic profile compromised both *in vitro* osteoblast differentiation and *in vivo* bone formation capacity **(Figure 3C-G)**. Notably, restoration of mitochondrial ATP production **(Figure 6B-F)** by use of the active metabolite of Sonlicromanol (KH183) rescued bone formation potential **(Figure 7D- G)**, implying that OXPHOS activity is flexible and can be targeted for enhancing bone formation. Glycolysis represents the main source of energy in undifferentiated osteoprogenitor cells (hBMSC)^4,5^. Here, hBMSC with genetically impaired OXPHOS exhibited the capacity to increase glycolysis to preserve ATP production, indicating compensatory bioenergetic flexibility in hBMSC **(Figure 1F-G)**. Nevertheless, the hBMSC cellular profile was compromised with a lower number of CFU-f, altered differentiation profile, and impaired *in vivo* bone formation capacity **(Figure 2)**. While bioenergetic flexibility in hBMSCs provides adequate ATP levels by increased glycolytic rates, further differentiation process to mature functional osteoblasts appears to depend to a larger degree on intact, highly active OXPHOS^5^. Therefore, impaired bone formation capacity of hBMSCs from m.3243A>G carriers could be explained by compromised OXPHOS.

In addition to decreased mitochondrial ATP production, compromised OXPHOS activity have secondary effects that can contribute to impaired bone formation. First, TCA cycle metabolites are important substrates for bone formation. Both citrate and alpha-ketoglutarate are metabolic substrates for amino acid biosynthesis, and alpha-ketoglutarate and succinate are cofactors for the biosynthesis of bone organic matrix components such as collagen type I^57^. The levels of these metabolites were lower in m.3243A>G hBMCS, as well as mitochondrial ATP production. While overall ATP production was maintained by bioenergetic flexibility towards a highly glycolytic profile **(Figure 1F-G)**, the compromised OXPHOS could impede the production of TCA cycle metabolites needed for bone formation with a negative effect on bone formation capacity **(Figure 3A-G)**. Second, impaired OXPHOS activity activates MSR^14,26^, and in line with this, MSR-related genes were upregulated in m.3243A>G carrierś hBMSCs **(Figure 2D)**. MSR consumes ATP for transcription of new genes, synthesis of new proteins and enzyme activity^14^. Therefore, although the total ATP production is preserved in carrierś hBMSC, ATP can be directed to increased MSR-related ATP consumption rather than bone formation. Collectively, secondary effects of impaired OXPHOS could have negative implications for the differentiation of osteoprogenitor cells and overall bone formation, either directly or indirectly. Future studies are needed to address these effects in maintaining bone homeostasis.

Restoring normal mitochondrial functions could improve the osteogenic potential of m.3243A>G hBMCS. Recently, Gao *et al*.^62^ demonstrated that suppression of MSR, specifically the mitochondrial unfolded protein response (UPR^mt^), enhanced mitochondrial function and osteogenic potential of human urine-derived stem cells from m.3243A>G carriers. Notably, our present treatment approach – using KH183 – specifically directed to improve the genetic impairment of OXPHOS, increased not only mitochondrial ATP production but also reduced glycolysis-associated lactate production, and increased TCA metabolites citrate, alpha-ketoglutaric acid and succinate **(Figure 6)**. In addition, the expression of MSR-related genes was reduced and all together, there was an upregulation of bone formation-related genes, such as *LRP5* and *RUNX2*, along with an increased mineralisation capacity **(Figure 7)**. This provides the proof-of-concept for OXPHOS as a gateway for bone formation, which has potential therapeutic implications.

While pre-clinical studies have documented the importance of intact mitochondrial function in bone homeostasis, the clinical significance remains to be detailed. Our previous study on a larger cohort of m.3243A>G carriers show lower BMD compared to controls unmatched on weight, also shown by Geng et al.^16,63^. In addition, epidemiological studies and case descriptions suggest individuals with inherited mitochondrial diseases have increased bone fragility^15,64,65^. Our results on human bone cells demonstrate that impaired OXPHOS impedes the differentiation of osteoprogenitors and their bone formation capacity. However, these effects were not reflected to the same degree in the skeletal phenotype of m.3243A>G carriers. While bone histomorphometry and μCT indicated that the m.3243A>G variant is associated with delayed bone formation and lower bone mass, the limited number of participants with this rare pathogenic variant and their lower BMI compared to the controls are a potential explanation for the differences observed **(Figure 5 and Table S4)**. Conversely, the discrepancy between the cellular and the clinical outcomes may also be explained by *in vivo* metabolic flexibility adapting to changes in energy substrates^66^, and cellular crosstalk with normal mitochondrial function cells, e.g. osteoclasts **(Figure 4)**. Thereby circumventing the development of significant skeletal phenotypes. This is supported by studies in mice where osteoprogenitors adapt to inhibition of mitochondrial respiration by increasing glycolysis and TCA metabolite succinate^6^.

Collectively, hBMSCs with genetically impaired OXPHOS exhibited a highly glycolytic profile and reduced bone formation potential. Furthermore, pharmacologically restoring OXPHOS function improved hBMSCs bone formation capacity. In conclusion, manipulating mitochondrial function – specifically OXPHOS – offers valuable insight into the metabolic regulation of osteogenesis and has the potential to enhance bone formation in conditions like osteoporosis that are characterized by inadequate bone formation.

## Limitations of the Study

This study demonstrates the importance of mitochondrial function for osteogenesis beyond providing ATP for cellular differentiation using unique samples from individuals carrying pathogenic variant m.3243A>G that led to impaired OXPHOS activity. Some limitations to be considered are common to cell culture studies. One shortcoming is that different types of bone cell are studied isolated, while the bone microenvironment is rich in different cell types, stages of differentiation and maturity, which may mask specific disease manifestations. Co-culturing different cell types could overcome this limitation. However, the relative contribution of each population to the different parameters analysed could not be discernible, especially in cells with highly different metabolic profiles like hBMSCs, osteoblasts, and osteoclasts. Another shortcoming is that we cultured the cells under standard culture conditions, such as on a plastic surface and atmospheric oxygen, while *in vivo* bone cells grow on the bone surface and under hypoxic conditions. Overall, these *in vitro* conditions may mask some of the *in vivo* effects. An intrinsic limitation of studying rare genetic conditions, such as m.3243A>G, is the limited accessibility to samples and heterogeneous disease presentation. However, the studies of rare diseases are important for modelling the basic biology of complex human diseases. To address this limitation, we included healthy individuals matched by sex-, age-, and BMI, which are major effectors of bone health, combined with an unusually high number of m.3243A>G carriers.

## Materials and Methods

### Participants and sample collection

Participants were subjects >18 years old that carry the pathogenic mitochondrial genetic variant m.3243A>G and healthy controls matched for sex, age, and BMI. Exclusion criteria: Renal (creatinine > 90 µmol/l) or liver (AST > 3 times the upper limit) dysfunction, medical treatment influencing bone metabolism (oral corticosteroid <12 weeks, anti-osteoporosis treatment, sex steroids, anti-convulsant), pregnancy, excessive consumption of alcohol (>14 unit/week), anticoagulants or pre-existing coagulopathy, allergy to lidocaine, morphine or diazepam.

This study is a cross-sectional and case-control study including two parts: **(A)** collection and study of bone biopsies and **(B)** collection of cells and analysis of cellular studies. *Part A. Bone studies.* Tetracycline-labelled bone biopsies were collected in 2016 from seven individuals carrying the m.3243A>G pathogenic variant and seven age- and sex-matched healthy individuals as controls. *Part B. Cellular studies.* Ten individuals carrying the m.3243A>G pathogenic variant participated in the cellular studies – five of them were also part of the bone study – with ten age-, sex-, and BMI-matched healthy individuals as controls. Bone marrow aspiration, blood samples and DXA scans were assessed in 2021 and 2022.

### Isolation of bone marrow mesenchymal stem cells from bone marrow aspirates

Bone marrow samples were obtained from the iliac crest by aspiration of 10-15 mL after infiltration of the area with local anaesthetic (10 mg/mL lidocaine) (Amgros). The samples were mixed 1:1 with Minimum Essential Media (MEM) (Gibco) containing heparin (100 U/mL) (Amgros). Low-density mononuclear cells were isolated through centrifugation with a Lymphoprep density gradient (density = 1.077 ± 0.001 g/cm^2^) (STEMCELL Technologies). hBMSCs were selected through the process of plastic adherence. Cells were cultured in MEM containing 10% FBS, 1% Penicillin/Streptomycin (P/S) (Gibco # 15140130), 1 mM Sodium

Pyruvate (Gibco # 11360039), 2 mM Glutamax (Gibco # 35050-038), 1% nonessential amino acids (Gibco # 11140050), and 50 µg/mL uridine (Sigma-Aldrich # U3003) incubated at 5% CO2 at 37°C. The media was changed for the first time after 7 days and every second day after that until 70% confluence (passage 0), when hBMSCs were cryopreserved and sub-cultured (passages 1, 2, and 3) for further studies.

### Culture of hBMSC

hBMSCs were sub-cultured in MEM containing 10% FBS, 1% P/S (Gibco # 15140130), 1 mM Sodium Pyruvate (Gibco # 11360039), 2 mM Glutamax (Gibco # 35050-038), 1% nonessential amino acids (Gibco # 11140050), and 50 µg/mL uridine (Sigma-Aldrich # U3003) in standard cell culture conditions (37°C, 85% humidity, and 5% CO2). hBMSCs were routinely tested for Mycoplasma and cultured in Mycoplasma-negative conditions. The medium was changed every other day.

### Cell count

The hBMSC were counted using via-1 cassettes in the automated cell counter NC-200 (Chemometec, Allerod, Denmark) according to the manufacturer’s instructions. Briefly, cell suspension was loaded in a via-1 cassette that contains DAPI as a nuclear staining for determining the non-viable cells concentration (Cnv). Another aliquot was mixed with cell lysis buffer and loaded in the via-1 cassette that contains DAPI as a nuclear staining for determining the total cell concentration (Ct). DAPI fluorescence was detected using peak excitation at 365 nm and emission at 410-460. Cell viability was calculated according to the following formula, viability = (Ct − Cnv)/Ct, and expressed as percentage.

### Sonlicromanol and KH183

Sonlicromanol (laboratory name KH176) was developed using Trolox as an initial hit compound and an iterative, medicinal chemistry phenotypic screening approach to improve drugability. The screening was based on common pathophysiological characteristics found in primary skin fibroblasts derived from patients with primary mitochondrial disease^55^. The deviations found, as compared to primary skin fibroblasts from control individuals, are increased ROS levels and increased redox stress sensitivity. During pharmaco- and toxicokinetic studies, a major metabolite with a quinone structure, referred to as KH183, was identified both in plasma and tissues of different animal species treated with KH176. The metabolite KH183 has been shown to have a pharmacological activity that is at least comparable to the activity of KH176.

### DNA isolation

DNA for heteroplasmy analysis was isolated from cultured hBMSC (passage 2 or 3) and mature osteoclasts (day 9 of differentiation). Cells were collected by trypsinization, washed with phosphate-buffered saline (PBS), centrifuged at 16,000 x g at 4°C, and stored at - 80°C before DNA isolation. DNA was isolated using Puregene Core Kit B (Qiagen) according to the manufactureŕs instructions.

### Mitochondrial DNA m.3243A>G heteroplasmy analysis

To quantify the fractional abundance of mitochondrial DNA with m.3243A>G, droplet digital PCR (ddPCR) was conducted. Each 22 µL ddPCR reaction consisted of 11 µl 2x ddPCR SuperMix for probes (no dUTP) (Bio-Rad, Hercules, California, USA), 909 nM of each primer, 568 nM of each probe, and 5 µl template DNA. Reactions were prepared in semi-skirted 96 well PCR plates (Bio-Rad, Hercules, California), and droplets were generated using an Automated Droplet Generator (Bio-Rad, Hercules, California). Following droplet generation, PCR plates were sealed with pierceable foil (Bio-Rad, Hercules, California) at 180°C for 5 seconds, and PCR was performed on a CFX96 Touch Deep Well Real-Time PCR System (Bio-Rad, Hercules, California). After optimisation of the assay, PCR conditions were 95°C for 10 minutes, 50 cycles of 94°C for 30 seconds and 56°C for 1 minute, and 98°C for 10 minutes with a ramp rate of 1°C/s. The plate was kept on hold at 4°C for a minimum of 30 minutes or at 12°C for a minimum of 4 hours, as described previously^67^, after which the plate was incubated for 10 minutes at room temperature and read on QX200 Droplet Reader (Bio-Rad, Hercules, California).

Every ddPCR analysis included a positive template control run in duplicates, a negative template control run in five wells, and a minimum of two no-template control (NTC) for a clean workflow quality control check. Only wells with a minimum of 10,000 accepted droplets were analysed, and thresholds were manually set at 1,200 in channel 1 (FAM) and 2,200 in channel 2 (HEX). Thresholds were defined during optimization using positive and negative controls with gating based on fluorescence amplitude in 1D and 2D plots. Data was analysed with QuantaSoft Analysis Pro 1.0 software (Bio-Rad, Hercules, California). A sample was scored positive for m.3243A>G if the concentration was above the false positive rate. The fractional abundance was calculated based on the concentration of both m.3243A>G and wild-type mitochondrial DNA for each sample.

### Bioenergetic analysis of hBMSC

Mitochondrial bioenergetics was performed using Seahorse XFe96 extracellular flux analyzer (Seahorse, Agilent, USA) to measure the Oxygen Consumption Rate (OCR) and Extracellular Acidification Rate (ECAR) simultaneously. hBMSC (passage 1) were seeded at 6,000 cells per well in 80 µL of MEM media (MEM 10% FBS + 1% P/S + 1 mM Sodium Pyruvate + 2 mM Glutamax + 1% nonessential amino acids + 50 µg/mL uridine) in a Seahorse 96-well cell culture plate (Agilent, USA). After 24 hours, the cells were treated for 72 hours with either vehicle (DMSO) or KH183 (5 µM) in cell culture media (MEM + 10% FBS + 1% P/S). An hour before the measurement, the cell culture medium was replaced with the assay medium and incubated in a non-CO2 environment at 37°C. The assay medium was prepared by supplementing Seahorse XF DMEM medium (Agilent, #103575-100) with 10mM glucose (Sigma-Aldrich, G8769), 1 mM Sodium Pyruvate (Gibco, 11360-039), and 2 mM Glutamine (Sigma-Aldrich, G7513).

For the analysis of mitochondrial respiration, we used the Mito Stress Test assay (Agilent, #103015-100) as previously described^68^. Two groups of sequential drug injections were used:**(1)** uncoupler FCCP (Carbonyl cyanide 4-(trifluoromethoxy) phenylhydrazone; 1 µM final/well), and inhibitors Rotenone/Antimycin A (0.5 µM final/well); **(2)** ATP-synthetase inhibitor Oligomycin (1 µM final/well), and Rotenone/Antimycin A (0.5 µM final/well). For the ATP production analysis, we used the second group of injections with Oligomycin and Rotenone/Antimycin A. For the glycolysis analysis, we used the Glycolytic Rate assay (Agilent ##103344-1) by injecting sequentially Rotenone/Antimycin A (0.5 µM final/well) and 2- deoxyglucose (50 mM final/well). The parameters obtained were calculated using the average of the three cycles measured according to the manufacturer’s recommendations.

After OCR and ECAR measurements were completed, we added Hoechst 33342 (Invitrogen, H3570) to each well as nuclear staining (20 µM/final well) for 15 minutes in the dark at 37°C. The cells were imaged by Cytation1 (Biotek, Agilent) using Cell Imaging software (Agilent) to count the number of cells in each well. The OCR and ECAR values were normalised to the Hoechst-stained cell counts, with the normalization unit set to 1,000 cells.

The normalized OCR and ECAR data were exported from Wave Software (Agilent) to Seahorse Analytics for the calculations as previously described^68^.

### Glucose uptake

The glucose uptake was determined by seeding hBMSCs (passage 3) in a 96-well black/clear bottom plate (Perkin Elmer) at a density of 6,000 cells/well (6 wells/group). After 24 hours, the cells were starved for 6 hours using low glucose (1 g/L) DMEM without FBS. Next, cells were incubated with glucose-free media without FBS with 10 µg/mL fluorescent 2-deoxyglucose (2- NBDG) (Caiman) for 2 hours. Finally, the cells were stained with Hoechst 33342 (Invitrogen) (1 µg/mL) for 10 min. The fluorescence was measured using the Cytation 1 Cell Imaging Multi- Mode Reader (BioTek Instruments, Agilent) using the DAPI channel for Hoechst 33342 (Excitation 377 nm, Emission 477 nm) and the GFP channel for 2-NBDG (Excitation 500/24 nm, Emission 542/27 nm). The mean fluorescent intensity of 2-NBDG was normalized to the number of Hoechst-stained cells.

### Mitochondrial Mass Determination with MitoTracker Green

Mitochondrial mass was determined with MitoTracker Green FM (Invitrogen, USA, M7514). We seeded 4,000 cells in a 96-well black/clear bottom plate (Perkin Elmer) (6 wells/group). After 24 hours, cells were washed with PBS and incubated with 100 nM MitoTracker Green and 20 µM Hoechst in HBSS (Hankś Balanced Salt Solution) for 30 minutes at 37°C. The fluorescence was measured using the Cytation 1 Cell Imaging Multi-Mode Reader (BioTek Instruments, Agilent) using the DAPI channel for Hoechst 33342 (377 nm/447 nm) and the GFP channel for MitoTracker Green (469 nm, 525 nm). The mean fluorescent intensity of MitoTracker Green was normalized to the number of Hoechst-stained cells.

### RNA isolation

hBMSC (passage 2) were seeded in 6-well plates at a density of 100,000/well (2 wells/group) in S-MEM media (MEM 10% FBS + 1% P/S + 1 mM Sodium Pyruvate + 2 mM Glutamax + 1% nonessential amino acids + 50 µg/mL uridine). After 24 hours, the cells were treated with either vehicle (DMSO) or KH183 (5 µM) in cell culture media (MEM + 10% FBS + 1% P/S) for 72 hours. Cells were washed once with PBS, and then 250 µL of TRIzol (Invitrogen #15596-018) was added. Then, the cells were detached using a cell scraper (VWR, #734-2602), collected in a pre-chilled microtube, and stored at -80°C.

RNA isolation was performed using 0.2 volumes of chloroform to allow phase separation. After 15 minutes of incubation, the suspension was centrifuged at 12,000 x g for 15 min at 4°C. The upper phase was transferred to a new 1.5 mL tube and kept on ice. Then, 0.6 volumes of 96% EtOH were added to the upper phase and vortexed. The suspension was then loaded onto an EconoSpin column (Epoch Life Science, Inc.) and centrifuged at 13,000g for 30 seconds. The column was washed three times with 500 µL RPE (80 % EtOH, 10 mM Tris pH 7.5) at RT with centrifugation at 13,000g for 30 seconds. Then, the column was centrifuged dry at 13,000g for 2 min. Afterwards, the column was incubated at RT for 5-10 minutes with the lid open to evaporate ethanol completely. Next, the RNA was eluted in 15 µL DEPC-treated H2O. The column was incubated for at least 2 minutes at RT and centrifuged at 16,000g for 1 minute at RT. The RNA concentration and quality were measured using the Thermo Scientific™ NanoDrop™ One Microvolume UV-Vis Spectrophotometers, and the RNA was stored at -80°C until sequencing.

### RNA sequencing

RNA sequencing was done at Aalborg University Hospital by a non-stranded and polyA-selected RNAsequencing library preparation. RNAseq library preparation was done using the TruSeq Stranded mRNA Library Prep Kit (Illumina) according to the manufactureŕs instructions. Sequencing is carried out as 2 × 150 bp paired-end on a NovaSeq 6000 (Illumina). Sequencing reads were mapped to the human genome (GRCh38) using MANE selected REfSeq genes^69^.

### Proteomics sample preparation

hBMSC (passage 2)(250,000 cells) were seeded in T25 flasks in MEM media (MEM 10% FBS + 1% P/S + 1 mM Sodium Pyruvate + 2 mM Glutamax + 1% nonessential amino acids + 50 µg/mL uridine). After 24 hours, the cells were treated with either a vehicle (DMSO) or KH183 (5 µM) in cell culture media (MEM + 10% FBS + 1% P/S) for 72 hours. Then, the cells were washed once with PBS, 1 mL of trypsin was added to the flasks and incubated for 3-5 min at 37°C. Samples were centrifuged at 200 x g for 5 minutes, and the supernatant was removed. The cell pellet was resuspended in 1 mL PBS and centrifuged at 16,000 g at 4°C. The supernatant was removed, and the cell pellet was resuspended in 1 mL PBS. The centrifugation step was repeated. The supernatant was discarded, and the pellet was stored at -80°C. Cell pellets were lysed in a HEPES lysis buffer (4% SDS (Invitrogen), 50 mM HEPES (Sigma Aldrich), and protease inhibitor (Roche, Complete mini), pH 7.4). Samples were further lysed by sonication in a water bath (Branson 2800 Ultrasonic Cleaner) for 10 seconds with 30 seconds of ice incubation between cycles (a total of 3 cycles). Sonicated lysates were centrifuged at 12,000 g for 10 minutes at 4°C. Protein concentrations of the supernatants were determined with the BCA protein assay (Thermo Scientific) according to the manufacturer’s instructions.

### Tandem Mass Tag Labeling

hBMSC samples were tagged with TMT10plex™ and TMTpro™16plex Isobaric Label Reagent Set (Thermo Scientific, #90111 and #A44520, respectively). From each sample, 100 µg protein was in-solution trypsin digested and TMT labelled according to the manufacturer’s instructions. Following the TMT labelling, all samples within each study were pooled. After a strong cation exchange (SCX) purification (Phenomenex, #8B-S010-EAK), peptides were loaded to an Immobiline™ DryStrip Gel (GE Healthcare, #11534985) isoelectric focusing (IEF) separation was performed. The IEF gels were then cut into 10 equal pieces, purified in PepClean C18 spin columns (Thermo Scientific, #11824141) according to the manufacturer’s instructions, and then vacuum centrifuged until dryness and stored at -20°C until nanoLC-MS/MS analyses.

### NanoLC-MS/MS and Proteomics Database Search

Nano-liquid chromatography tandem-mass spectrometry (nanoLC-MS/MS) was performed on an EASY nanoLC-1200 coupled to a Q-Exactive™ HF-X Quadrupole-Orbitrap™ Mass Spectrometer (Thermo Scientific) as previously described^70^. Briefly, MS was operated in positive mode using pre- and analytical columns: acclaim PepMap 100, 75 µm × 2 cm (Thermo Scientific) and EASY-Spray PepMap RSLC C18, 2µm, 100 AÅ, 75 µM × 25 cm (Thermo Scientific), respectively, to trap and separate peptides in a 170-minute gradient with 5-40 % acetonitrile, 0.1 % formic acid. Higher-energy collisional dissociation (HCD) was used for peptide fragmentation, and the normalized collision energy (NCE) was 35. Full scan (MS1) and fragment scan resolutions were set at 60,000 and 45,000, respectively. Automatic gain control (AGC) for MS1 and MS2 were set at 1×10^6^ with a scan range between 372-1,500 m/z and at 1×10^5^ with a fixed first mass of 110 m/z, respectively. Up to 12 of the most intense peaks of the full scan were fragmented with data-dependent analysis (DDA). Unassigned and +1 charge states were excluded from fragmentation, and the dynamic exclusion duration was 15 seconds. Each fraction was LC-MS/MS analyzed twice, where peptides identified from >9 scans in the first analysis were excluded from fragmentation in the second analysis.

### Proteomics Database Search

All LC-MS/MS results were merged and submitted for the database search for identification and quantification of proteins using Proteome Discoverer 3.0 (Thermo Scientific). 20,401 reviewed *Homo sapiens* Uniprot sequences were used as reference proteome (Swiss-Prot; downloaded on 02.12.2022) using the Sequest algorithm. Precursor mass tolerance was 10 ppm, and fragment mass tolerance was 20 mmu. The maximum number of allowed missed cleavages was two. Oxidation of methionine was set as dynamic modification, and static modifications were carbamidomethylation of cysteines and TMT-plex-labels on lysine and peptide N-terminus. The co-isolation threshold was set at 35%, and the identification false discovery rate was 0.01 at both peptide and protein levels. Proteins with more than 2 quantitative peptide scans and at least one unique peptide were considered as quantified and were included in the further proteomics data analysis. Criteria for the differentially expressed proteins were set at p < 0.05 and |FC|>1.2 for protein abundance.

### Colony forming units-fibroblast (CFU-f) assay

For the assessment of CFU-f, hBMSC on the day of isolation were plated, 0.5 and 1 million cells in T25 flasks with MEM 10% FBS + 1% P/S + 1 mM Sodium Pyruvate + 2 mM Glutamax + 1% nonessential amino acids + 50 µg/mL uridine. The media was renewed on days 7 and 10. After 14 days, colonies displaying more than 50 cells were visually counted after Crystal Violet staining (Sigma-Aldrich).

### Metabolomics sample preparation

hBMSC (passage 1) were seeded in 6-well plates at 100,000 cells/well (2 wells/group) in media (MEM 10% FBS + 1% P/S + 1 mM Sodium Pyruvate + 2 mM Glutamax + 1% nonessential amino acids + 50 µg/mL uridine). After 24 hours, the cells were treated with either vehicle (DMSO) or KH183 (5 µM) in cell culture media (MEM + 10% FBS + 1% P/S) for 72 hours. Cells were washed once with warm PBS and incubated for 90 min at 37°C with MEM containing 2.5 mM of [U- ^13^C]glucose. After incubation, the medium was collected, and the cells were washed with cold PBS (4°C) to stop metabolic reactions. The cells were lysed, and metabolites were extracted with 70% cold ethanol. The extract was centrifuged at 20,000g for 20 min at 4°C to separate soluble and insoluble components. Pellets were dissolved in KOH (1 M) at room temperature and analyzed for protein content by the BCA assay (ThermoFisher) according to the manufacturer’s instructions.

### Metabolic mapping using gas chromatography coupled to mass spectrometry (GC–MS)

^13^C-enrichment of metabolites was determined using GC-MS as previously described^71^. Briefly, extracts were reconstituted in water, acidified, and metabolites were extracted into an organic phase with 96% ethanol/benzene and derivatized using N-tert-butyldimethylsilyl-N- methyltrifluoroacetamide. The relative abundance of ions based on their mass-to-charge (m/z) values was determined as previously described^71^. Natural ^13^C-abundance was corrected and calculated as previously described^72^. Data are presented as a percentage of labelling of the isotopologue M + X, where M corresponds to the molecular weight of the unlabelled molecule and X is the number of ^13^C-enriched carbon atoms in the molecule^73^. The TCA cycle metabolites were analyzed as the result of the first turn of the cycle with the addition of only two ^13^C- enriched carbon atoms in the molecule (M + 2) and the second turn with the addition of four ^13^C-enriched carbon atoms in the molecule (M + 4).

### Cell Viability

Cellular viability was determined by adding the CellTiter-Blue solution (Promega, Madison, USA) for 1 hour according to the manufacturer’s instructions. The absorbance was measured at a wavelength of 570 nm in a FLUOmega plate reader.

### Osteoblasts differentiation

hBMSCs (passage 1) were plated at a density of 20,000 cells/cm^2^ in MEM supplemented with 10% FBS and 1% P/S. Osteoblastogenesis was induced on the following day by media supplemented with 10 nM b-glycerophosphate (Sigma-Aldrich), 10 nM dexamethasone (Sigma-Aldrich), 50 μg/mL vitamin C (Sigma-Aldrich) and 50 μg/mL vitamin D (Sigma-Aldrich). The medium was changed every other day for the duration of the differentiation.

### Alkaline phosphatase (ALP) activity assay

ALP activity was analysed after 7 days of osteoblast differentiation in 96-well plates (6 wells/group). The cell number (viable cells) was determined as described before. Subsequently, the cells were rinsed with TBS (20 mM Trizma base (Sigma-Aldrich), 150 mM NaCl (Thermo Fisher), pH = 7.5) and fixed in 3.7% formaldehyde-90% ethanol (Sigma-Aldrich) for 30 seconds at room temperature. A reaction mixture containing 50 mM NaHCO3, 1 mM MgCl2 (Sigma-Aldrich), and 1 mg/ml of p-nitrophenyl phosphate (Sigma-Aldrich) was added into each well and incubated at 37°C for 20 minutes. The reaction was stopped by adding 50 μL of 3 M NaOH. Absorbance was measured at 405 nm in a FLUOmega plate reader. ALP enzymatic activity was normalized to cellular viability.

### Alizarin Red Staining

hBMSCs (passage 1) were seeded in 4-well plates. The day after, cells were induced to differentiate into osteoblasts using osteoblastic induction medium. On day 14 of osteoblast differentiation, mineralized matrix formation was measured using Alizarin Red staining (AR- S). Cells were fixed with 70% ice-cold ethanol for at least 1 hour at -20°C before adding AR-S (40 mM; Sigma-Aldrich) for 10 minutes at room temperature (RT). All samples were stained and quantified at the same time. The level of calcium deposition was quantified by elution of AR-S. The absorbance of the eluted dye was assessed at 570 nm in a FLUOmega plate reader.

### *In vivo* heterotropic bone formation assay

The *in vivo* heterotropic bone formation assay is used to evaluate the *in vivo* bone formation capacity of hBMSC (passage 2) by implanting them subcutaneously in immunodeficient mice as previously described^74^. Briefly, before implantation, we seeded 500,000 cells onto scaffold granules in 200 µL of cell culture medium (MEM + 10% FBS + 1% P/S) in cut-off syringes and incubated them overnight at 37°C and 5% CO2. The scaffold granules were Hydroxyapatite/tricalcium phosphate (HA/TCP) granules (Zimmer Scandinavia, Hørsholm, Denmark). Each mouse received two implants from the same participant. The mice were 8- week-old females NOD/SCID (NOD/LtSz-Prkdcscid).

### Adipocyte differentiation

hBMSCs (passage 1) plated at 30.000 cells/cm^2^ density. Adipogenesis was induced on the following day by DMEM supplemented with 10% FBS, 1% P/S, 100 nM dexamethasone (Sigma-Aldrich), 0.5 mM 3-isobutyl-1-methylxanthine (IBMX) (Sigma-Aldrich), 1 µM BRL (Rosiglitazone) (Sigma-Aldrich), 2 µg/mL insulin (Sigma-Aldrich). The medium was changed every other day for the duration of the differentiation.

### Lipid droplets quantification

Lipid droplet quantification was used to evaluate adipocyte differentiation, which was determined with Nile Red (Sigma-Aldrich) staining. We seeded cells (6 wells/group) in a 96-well black/clear bottom plate (Perkin Elmer). After 24 hours, cells were induced to differentiate into adipocytes using adipogenic induction medium. On day 7, cells were washed with PBS and incubated with Nile Red (5 µg/mL) for 15 minutes in the dark at 37°C. The fluorescence was quantified using FLUOmega plate reader at 485 nm excitation and 572 nm emission. The Nile Red fluorescent was normalized to cellular viability per well.

### Osteoclasts differentiation

Peripheral Blood Mononuclear Cells (PBMCs) were isolated from 50 mL of EDTA blood by differential centrifugation using Ficoll-Paque (Cytiva) as a density gradient. The PBMCs were washed twice with PBS (Thermo Fisher) before counting the cells with trypan blue (Thermo Fisher) using an automated cell counter – Countess (Invitrogen). Cells were seeded at a density of 5×10^7^ in T75 or 1.7×10^7^ in T25 culture flasks and differentiated to mature osteoclasts over 9 days in alpha-MEM (Thermo Fisher) with 10% FBS (Thermo Fisher) and 1% P/S (100 U/mL penicillin (GIBCO), and 100 μg/mL streptomycin (GIBCO) (Invitrogen). For the first 2 days, the cells were induced with 25 ng/mL M-CSF (R&D Systems, Abingdon, UK), after cells were induced with 25 ng/mL M-CSF and 25 ng/mL RANKL (R&D Systems, Abingdon, UK). After 9 days of maturation, 12 systematic and evenly distributed pictures were taken to quantify the number of nuclei per osteoclast and the number of osteoclasts with two or more nuclei using a ckx41 microscope with an SC30 camera (Olympus Corporation, Tokyo, Japan).

### Bone resorption assay

The osteoclasts bone resorptive activity were performed as previously described^50^. Briefly, matured osteoclasts (day 9 of induction) were detached by accutase (PSS, Pasching, Austria) and were seeded onto bone slices (50,000 cells/bone slice) (www.boneslices.com, Jelling, Denmark) in eight different wells of a 96-well plate. The cells were cultivated for 72 hours in alpha-MEM with 10% FBS, 1% P/S, 25 ng/mL M-CSF, and 25 ng/mL RANKL. The media was stored at -20 °C for later measurements of TRAcP activity. The bone slices were washed with water and polished with a cotton swap to remove the cells. The cleaned bone slices were subsequently stained with toluidine blue (Thermo Fisher). Bone resorption was quantified by assessing the percentage of eroded surface per bone surface according to the occurrence of two types of resorption patterns: pits and trenches^50^. A pit was defined as an excavation, circular in appearance, with well-defined edges, and where the ratio between the length and the width of the excavation did not exceed two. A trench was defined as an elongated and continuous excavation with well-defined edges that were at least two times longer than its width. The analysis was performed blinded and randomized.

### TRAcP activity analysis

TRAcP activity was analysed in frozen cell culture media collected during the differentiation and bone resorption assays. In brief, 10 µL of cell culture media was analysed in duplicate by incubation with TRAcP reaction buffer (1 M acetate (Sigma-Aldrich), 0.5% Triton X-100 (Sigma-Aldrich), 1 M NaCl (Sigma Aldrich), 10 mM EDTA (VWR), pH 5.5), 50 mM L-Ascorbic acid (Sigma-Aldrich), 0.2 M disodium tartrate (Sigma-Aldrich), and 82 mM 4-nitrophenylphosphate (Sigma-Aldrich), for 15 minutes at 37 °C in the dark. The reaction was stopped by adding 100 µL of 0.3 M sodium hydroxide (VWR). The absorbance was measured at 400 nm using a microplate reader (Synergy HT, Biotek).

### Bioenergetic analysis of mature osteoclasts

The matured osteoclasts (day 8 of induction) were detached by accutase (Biowest) and seeded in seahorse cell culture plates (40,000 cells/well) (Agilent) in 6-8 wells/parameter. The cells were incubated in alpha-MEM with 10% FBS, 1% P/S, 25 ng/mL M-CSF, and 25 ng/mL RANKL for 24 hours before the assay, as previously described^75^. Briefly, the cells were washed with Seahorse media (non-buffered DMEM (Agilent # 103575-100) supplemented with 1 mM glucose (Agilent), 1 mM sodium pyruvate (Agilent), and 2 mM glutamine (Agilent)), 180 µL of Seahorse media was added to each well, and the plate was incubated in a non-CO2 incubator for 60 min.

For the analysis of mitochondrial respiration, we used the Mito Stress Test assay (Agilent, #103015-100) as previously described^68^. Briefly, two groups of sequential drug injections were used: **(1)** uncoupler FCCP (Carbonyl cyanide 4-(trifluoromethoxy) phenylhydrazone; 2.5 µM final/well), and inhibitors Rotenone/Antimycin A (0.5 µM final/well); **(2)** ATP-synthetase inhibitor Oligomycin (1.5 µM final/well), and Rotenone/Antimycin A (0.5 µM final/well). For the ATP production analysis, we used the second group of injections with Oligomycin and Rotenone/Antimycin A. For the glycolysis analysis, we used the Glycolytic Rate assay (Agilent #103344-100) by injecting sequentially Rotenone/Antimycin A (0.5 µM final/well) and 2-deoxyglucose (50 mM final/well). The parameters obtained were calculated using the average of the three cycles measured according to the manufacturer’s recommendations.

### Bone biopsies

Transiliac crest bone biopsies were collected using a modified Bordier procedure after each study subject underwent double labelling with tetracycline administrated orally. Before the bone biopsy, the participants ingested tetracycline hydrochloride 250 mg three times daily on days 1-3 and days 13-15. On day 20, a 7 mm diameter core was obtained across the iliac crest. The specimens were subsequently fixed and stored in 70% ethanol/30% water at 4° C until dehydrated and embedded non-decalcified in methyl-methacrylate. A Jung Model K microtome was used to cut 7-µm–thick sections for detailed bone and adipocyte histomorphometric analyses.

### Micro-CT scanning

The whole transiliac bone biopsy specimens were scanned using a µCT scanner (µCT 50, Scanco Medical AG, Brüttisellen, Switzerland) with an isotropic voxel size of 6 µm (X-ray tube: 155 µA, 90 kVp, integration time 1500 ms) to quantify the 3D microarchitectural properties of the cancellous bone^76^. All specimens were scanned in the same orientation. The 3D-image data sets, filtered with a Gaussian filter (sigma = 0.8, support = 1) and segmented with an optimal threshold of 180, consisted of approximately 600 consecutive 16-bit grey scale images. These images were used to quantify (after segmentation with a fixed optimal threshold) bone volume fraction (BV/TV, %), trabecular thickness (Tb.Th, µm), trabecular spacing (Tb.Sp, µm), trabecular number (Tb.N, mm^-1^), structure model index (SMI), connectivity density (mm^-3^), trabecular bone density (TBD, µg/cm^3^), cortical thickness (Ct.Th, mm) and cortical porosity (Ct.Po, %)^77^.

### Bone histomorphometry

The transiliac bone biopsy specimens were fixed in 70 % Ethanol and undecalcified embedded methylmethacrylate (MMA), before µCT scanned and sectioned for the bone histomorphometry on a SM2500 heavy-duty microtome (Leica). Central seven-µm-thick sections were either Masson’s trichrome stained for static histomorphometry or unstained for dynamic histomorphometry of tetracycline (mineralization) label and imaged on a VS200 slide scanner (Olympus)^78^. The slide scans were performed using a combination of bright-field and polarized light microscopy on the Masson-stained sections, and fluorescence microscopy of the tetracycline double-labels on the unstained sections, and manually analysed using the VS200 desktop software (Olympus). The static histomorphometry measures included osteoid surface per bone surface (OS/BS, %) and eroded surfaces per bone surface (ES/BS, %), where the eroded surfaces were divided according to whether they had neighbouring osteoid surfaces [(ES with neighbouring OS/BS, %) and (ES without neighbouring OS/BS, %)]^79^. The dynamic histomorphometry measures included the single and double labelled perimeter (sL.S/Pm and dL.S/Pm, µm) derived into the MS/BS (%) = M.Pm/B.Pm = (1/2 x sL.Pm + dL.S/Pm)/B.Pm, and mean interlabel width (Il.W, µm) derived into the mineral apposition rate (MAR, µm/day) = Il.W x Il.time (days) x π/4.

### Adipocyte histomorphometry

The bone marrow adiposity was investigated in scans of Masson’s trichrome stained sections, using point- and box-grids as previously described^80^. The point-grid measures allow us to estimate the adiposity area per marrow area (Ad.Ar/M.Ar, %), while the box-grid measures allow us to estimate the adipocyte density as the adipocyte profile number per marrow area (Ad.Pf.N/M.Ar), as well as the mean adipocyte profile diameter (Ad.Pf.Dm).

### Dual-energy X-ray absorptiometry (DXA) scan

Areal bone mineral density (aBMD) was measured at the lumbar spine (L1-L4), total hip and femoral neck using DXA (Hologic Discovery, Waltham, Massachusetts, USA). Z-scores and T-scores were calculated using the reference range provided by the manufacturer and the Third National Health and Nutrition Examination Survey reference^81^.

### Bioinformatics and Enrichment Analysis

RNA sequencing data analysis was performed in R as previously described^82^. Briefly, Differential gene expression was analyzed using DESeq2^83^ with a factor design (design = ∼ Match + Group), which regresses m.3243A>G carriers and controls variation to determine the differentially expressed (DE) genes. Linear variable (design = ∼ Match + hBMSC heteroplasmy) was used for each individual to determine the effect of heteroplasmy levels on the DE genes. DE genes were selected based on an adjusted *p*-value < 0.05. Pathway analysis was performed using Gene Ontology (GO) analyses using GOseqseq^84^ and EnrichR^85,86^. Mitochondrial proteins and their encoded genes were selected using Human MitoCarta3.0^21^. Enrichment analysis of the DE proteins (*p*-value <0.05) was performed using the gene name for each protein for the Reactome analysis on EnrichR^85,86^. The figures represent the main GO term found in the analysis, the children GO term were not shown in the figures.

## Statistics

The statistical significance was determined by a Wilcoxon signed-ranked test, a paired non-parametric test, unless stated otherwise. A *p*-value < 0.05 was considered significant (* *p* < 0.05, ** *p* < 0.01 and *** *p* < 0.001). The n values correspond to the number of individuals analysed in each experiment, the number of technical replicates is specified in each section of the materials and methods. The indicated statistical tests and graphs were performed using R version 4.1.1. Data was processed using tidyverse (v.1.3.2) and rstatix (v.0.7.1) for statistical analysis. The data was visualized with R packages ggplot2 (v.3.4.2) and ggpubr (v.0.4.0).

## Ethics Statement

Signed informed consent was obtained from all participants. This study conformed to the Declaration of Helsinki and was approved by The Ethic Committee of the Region of Southern Denmark (S-20100112, nr 46798, and S-20180170) and the Danish Civil Data Registry (j.nr. 2011-41-6172 and 19/7854) for the studies of bone biopsies and cellular samples, respectively. The heterotopic bone implant experiments were approved by The Danish Animal Experiments Inspectorate (License: 2017-15-0201-01210). ClinicalTrials.gov ID: NCT05483738.

## Data availability

Foldchange data of the RNASeq and proteomics studies are comprised in the excel sheets of Supplemental Table S1.

Due to data protection and study participant confidentiality, raw RNA seq and proteomics data is not publicly available. Any additional information required to reanalyze the data reported in this paper is available from the lead contact (Paula Fernandez Guerra upon request in an anonymized way.

## Acknowledgments

We thank Lotte Hørlyck, Charlotte Ejersted, and Julie Bjerrelund from the Department of Endocrinology, Odense University Hospital, for their assistance in collecting samples from the participants and analysing clinical data. We thank Tina K. Nielsen, Nicholas Ditzel, and Ahmed Sayed from The Molecular Endocrinology & Stem Cell Research Unit (KMEB), Molecular Endocrinology, Odense University Hospital (OUH), Denmark for their technical assistance in the isolation of bone marrow mesenchymal stem cells, heterotrophic bone formation assays, and analysis of osteoblast differentiation. Furthermore, we thank Jacob Bastholm Olesen, Clinical Cell Biology, Dept. of Pathology, OUH, for his technical assistance in the osteoclast assays. We also thank Kaja Søndergaard Laursen from the Department of Forensic Medicine at Aarhus University for her technical assistance in the histology analysis. The RNA sequencing analysis was performed at Dept. of Molecular Diagnostics, Aalborg University Hospital, with the assistance of Mads Sønderkær. We also thank Alexander Rauch for his advice in the analysis of RNA sequencing. The proteomics analysis was performed at the Research Unit of Molecular Medicine, Aarhus University, with the assistance of Margrethe Kjeldsen. The dynamic metabolic mapping was performed at the Department of Drug Design and Pharmacology, University of Copenhagen, with the assistance of Sebastian Weber Nielsen.

This work was supported by the Novo Nordisk Foundation (NFF18OC0052491 to ALF and NNF19OC0055047 to MF), Region of Southern Denmark (J.nr.18/51831 to ALF), and Entrepreneur Marius Pedersen Foundation (to ALF and PFG).

## Author contributions

P.F.G., A.L.F., and M.F. conceived the study. P.F.G. designed, performed and analysed most of the experiments. P.F.G. and P.K.K. performed and analyzed the respirometry assays, heterotrophic bone formation assay, and RNA sequencing experiments. S.K.T. performed and analyzed the droplet digital PCR for heteroplasmy analysis. P.F.G., P.K.K., and B.A. performed and analyzed the dynamic metabolic mapping experiments. P.H.A. included participants in the study and performed clinical evaluations. P.F.G., P.K.K. and J.P. performed and analyzed the data from the proteomic studies. P.F.G., P.K.K. and K.S. designed and analyzed the osteoclast experiments. P.F.G., A.L.F., and T.L.A. designed and analyzed the bone biopsy experiments. J.S.T. performed and analyzed the μCT of the bone biopsies. H.R and J.S kindly provided the KH183 compound and contributed with scientific advice. M.K. contributed to the interpretation and provided critical scientific advice. P.F.G., A.F.L., and M.F. wrote and revised the manuscript. All authors contributed to data interpretation, read, edited, and approved the final version of the manuscript.

## References

1. Delaisse, J.-M. et al. Re-thinking the bone remodeling cycle mechanism and the origin of bone loss. Bone 141, 115628 (2020).

2. Karsenty, G. & Khosla, S. The crosstalk between bone remodeling and energy metabolism: A translational perspective. Cell Metab. 34, 805–817 (2022).

3. Paula, F. J. A. de & Rosen, C. J. Bone Remodeling and Energy Metabolism: New Perspectives. Bone Res. 1, 72–84 (2013).

4. Lu, W., Duan, Y., Li, K., Qiu, J. & Cheng, Z. Glucose uptake and distribution across the human skeleton using state-of-the-art total-body PET/CT. Bone Res. 11, 36 (2023).

5. Sautchuk, R. & Eliseev, R. A. Cell energy metabolism and bone formation. Bone Rep. 16, 101594 (2022).

6. Tournaire, G. et al. Skeletal progenitors preserve proliferation and self-renewal upon inhibition of mitochondrial respiration by rerouting the TCA cycle. Cell Rep. 40, 111105 (2022).

7. Hollenberg, A. M., Smith, C. O., Shum, L. C., Awad, H. & Eliseev, R. A. Lactate Dehydrogenase Inhibition With Oxamate Exerts Bone Anabolic Effect. J. Bone Miner. Res. 35, 2432–2443 (2020).

8. Lin, C. et al. Impaired mitochondrial oxidative metabolism in skeletal progenitor cells leads to musculoskeletal disintegration. Nat. Commun. 13, 6869 (2022).

9. Trifunovic, A. et al. Premature ageing in mice expressing defective mitochondrial DNA polymerase. Nature 429, 417–423 (2004).

10. Jin, Z., Wei, W., Yang, M., Du, Y. & Wan, Y. Mitochondrial Complex I Activity Suppresses Inflammation and Enhances Bone Resorption by Shifting Macrophage-Osteoclast Polarization. Cell Metabolism 20, 483–498 (2014).

11. Picard, M. & Shirihai, O. S. Mitochondrial signal transduction. Cell Metab 34, 1620–1653 (2022).

12. Monzel, A. S., Enríquez, J. A. & Picard, M. Multifaceted mitochondria: moving mitochondrial science beyond function and dysfunction. Nat. Metab. 5, 546–562 (2023).

13. Martínez-Reyes, I. & Chandel, N. S. Mitochondrial TCA cycle metabolites control physiology and disease. Nat. Commun. 11, 102 (2020).

14. Monzel, A. S., Levin, M. & Picard, M. The energetics of cellular life transitions. Life Metab. 3, load051 (2023).

15. Gandhi, S. S., Muraresku, C., McCormick, E. M., Falk, M. J. & McCormack, S. E. Risk factors for poor bone health in primary mitochondrial disease. Journal of Inherited Metabolic Disease 40, 673–683 (2017).

16. Langdahl, J. H. et al. Mitochondrial Point Mutation m.3243A>G Associates With Lower Bone Mineral Density, Thinner Cortices, and Reduced Bone Strength: A Case-Control Study. Journal of Bone and Mineral Research 32, 2041–2048 (2017).

17. Jeppesen, T. D., Orngreen, M. C., Hall, G. V. & Vissing, J. Lactate metabolism during exercise in patients with mitochondrial myopathy. Neuromuscul. Disord. 23, 629–636 (2013).

18. Parikh, S. et al. Diagnosis and management of mitochondrial disease: a consensus statement from the Mitochondrial Medicine Society. Genet. Med. 17, 689–701 (2015).

19. Stewart, J. B. & Chinnery, P. F. The dynamics of mitochondrial DNA heteroplasmy: implications for human health and disease. Nat. Rev. Genet. 16, 530–542 (2015).

20. Frederiksen, A. L. et al. High Prevalence of Impaired Glucose Homeostasis and Myopathy in Asymptomatic and Oligosymptomatic 3243A>G Mitochondrial DNA Mutation-Positive Subjects. J. Clin. Endocrinol. Metab. 94, 2872–2879 (2009).

21. Rath, S. et al. MitoCarta3.0: an updated mitochondrial proteome now with sub-organelle localization and pathway annotations. Nucleic Acids Res. 49, gkaa1011- (2020).

22. Sturm, G. et al. OxPhos defects cause hypermetabolism and reduce lifespan in cells and in patients with mitochondrial diseases. *Commun*. Biol. 6, 22 (2023).

23. Lawton, A., Tripodi, N. & Feehan, J. Running on empty: Exploring stem cell exhaustion in geriatric musculoskeletal disease. Maturitas 188, 108066 (2024).

24. Picard, M. et al. Progressive increase in mtDNA 3243A>G heteroplasmy causes abrupt transcriptional reprogramming. Proc. Natl. Acad. Sci. 111, E4033–E4042 (2014).

25. Alston, C. L., Rocha, M. C., Lax, N. Z., Turnbull, D. M. & Taylor, R. W. The genetics and pathology of mitochondrial disease. J. Pathol. 241, 236–250 (2017).

26. Mick, E. et al. Distinct mitochondrial defects trigger the integrated stress response depending on the metabolic state of the cell. eLife 9, e49178 (2020).

27. Chung, H. K. et al. Growth differentiation factor 15 is a myomitokine governing systemic energy homeostasis. J. Cell Biol. 216, 149–165 (2017).

28. Poulsen, N. S. et al. Growth and differentiation factor 15 as a biomarker for mitochondrial myopathy. Mitochondrion 50, 35–41 (2020).

29. Li, Y. et al. Circulating FGF21 and GDF15 as Biomarkers for Screening, Diagnosis, and Severity Assessment of Primary Mitochondrial Disorders in Children. Front. Pediatr. 10, 851534 (2022).

30. Maresca, A. et al. Expanding and validating the biomarkers for mitochondrial diseases. J. Mol. Med. 98, 1467–1478 (2020).

31. Winter, J. M., Yadav, T. & Rutter, J. Stressed to death: Mitochondrial stress responses connect respiration and apoptosis in cancer. Mol. Cell 82, 3321–3332 (2022).

32. Barilani, M. et al. Age-related changes in the energy of human mesenchymal stem cells. J. Cell. Physiol. 237, 1753–1767 (2022).

33. Zhang, H., Menzies, K. J. & Auwerx, J. The role of mitochondria in stem cell fate and aging. Development 145, dev143420 (2018).

34. Yan, C. et al. Mitochondrial quality control and its role in osteoporosis. Front. Endocrinol. 14, 1077058 (2023).

35. Rauch, A. et al. Osteogenesis depends on commissioning of a network of stem cell transcription factors that act as repressors of adipogenesis. Nat Genet 51, 716–727 (2019).

36. Kegelman, C. D. et al. Skeletal cell YAP and TAZ combinatorially promote bone development. FASEB J. 32, 2706–2721 (2018).

37. Byun, M. R. et al. Canonical Wnt signalling activates TAZ through PP1A during osteogenic differentiation. Cell Death Differ. 21, 854–863 (2014).

38. Zou, M.-L. et al. The Smad Dependent TGF-β and BMP Signaling Pathway in Bone Remodeling and Therapies. Front. Mol. Biosci. 8, 593310 (2021).

39. Wu, M., Chen, G. & Li, Y.-P. TGF-β and BMP signaling in osteoblast, skeletal development, and bone formation, homeostasis and disease. Bone Res. 4, 16009 (2016).

40. Angelozzi, M., Karvande, A. & Lefebvre, V. SOXC are critical regulators of adult bone mass. Nat. Commun. 15, 2956 (2024).

41. Gadi, J. et al. The Transcription Factor Protein Sox11 Enhances Early Osteoblast Differentiation by Facilitating Proliferation and the Survival of Mesenchymal and Osteoblast Progenitors*. J. Biol. Chem. 288, 25400–25413 (2013).

42. Pan, J.-X. et al. YAP promotes osteogenesis and suppresses adipogenic differentiation by regulating β-catenin signaling. Bone Res. 6, 18 (2018).

43. Ambrosi, T. H. et al. Adipocyte Accumulation in the Bone Marrow during Obesity and Aging Impairs Stem Cell-Based Hematopoietic and Bone Regeneration. Cell Stem Cell 20, 771–784.e6 (2017).

44. Infante, A. & Rodríguez, C. I. Osteogenesis and aging: lessons from mesenchymal stem cells. Stem Cell Res. Ther. 9, 244 (2018).

45. Smith, R. L., Soeters, M. R., Wüst, R. C. I. & Houtkooper, R. H. Metabolic Flexibility as an Adaptation to Energy Resources and Requirements in Health and Disease. Endocr. Rev. 39, 489– 517 (2018).

46. Simonsen, J. L. et al. Telomerase expression extends the proliferative life-span and maintains the osteogenic potential of human bone marrow stromal cells. Nat Biotechnol 20, 592–596 (2002).

47. Lemma, S. et al. Energy metabolism in osteoclast formation and activity. Int J Biochem Cell Biology 79, 168–180 (2016).

48. Langdahl, J. H. et al. Lecocytes mutation load declines with age in carriers of the m.3243A>G mutation: A 10-year Prospective Cohort. Clin. Genet. 93, 925–928 (2018).

49. Grady, J. P., et al. mtDNA heteroplasmy level and copy number indicate disease burden in m.3243A>G mitochondrial disease. EMBO Mol. Med. 10, e8262 (2018).

50. Merrild, D. M. et al. Pit- and trench-forming osteoclasts: a distinction that matters. Bone Res. 3, 15032 (2015).

51. Halleen, J. M. Tartrate-resistant acid phosphatase 5B is a specific and sensitive marker of bone resorption. Anticancer research 23, 1027—1029 (2003).

52. Dobson, P. F. et al. Mitochondrial dysfunction impairs osteogenesis, increases osteoclast activity, and accelerates age related bone loss. Sci. Rep. 10, 11643 (2020).

53. Maclaine, K. D., Stebbings, K. A., Llano, D. A. & Havird, J. C. The mtDNA mutation spectrum in the PolG mutator mouse reveals germline and somatic selection. *BMC Genom*. Data 22, 52 (2021).

54. Smeitink, J. et al. Phase 2b program with sonlicromanol in patients with mitochondrial disease due to m.3243A>G mutation. Brain awae 277 (2024) doi:10.1093/brain/awae277.

55. Beyrath, J. et al. KH176 Safeguards Mitochondrial Diseased Cells from Redox Stress-Induced Cell Death by Interacting with the Thioredoxin System/Peroxiredoxin Enzyme Machinery. Sci Rep-uk 8, 6577 (2018).

56. Fernández-Vizarra, E. et al. Two independent respiratory chains adapt OXPHOS performance to glycolytic switch. Cell Metab. 34, 1792–1808.e6 (2022).

57. Viguet-Carrin, S., Garnero, P. & Delmas, P. D. The role of collagen in bone strength. Osteoporos. Int. 17, 319–336 (2006).

58. Riddle, R. C. et al. Lrp5 and Lrp6 Exert Overlapping Functions in Osteoblasts during Postnatal Bone Acquisition. PLoS ONE 8, e63323 (2013).

59. Vlashi, R., Zhang, X., Wu, M. & Chen, G. Wnt signaling: Essential roles in osteoblast differentiation, bone metabolism and therapeutic implications for bone and skeletal disorders. Genes Dis. 10, 1291–1317 (2023).

60. Ouwens, D. M. et al. Canonical WNT pathway inhibition reduces ATP synthesis rates in glioblastoma stem cells. Front. Biosci.-Landmark 27, 1 (2022).

61. Costa, R. et al. Impaired Mitochondrial ATP Production Downregulates Wnt Signaling via ER Stress Induction. Cell Rep. 28, 1949–1960.e6 (2019).

62. Gao, X. et al. ATF5, a putative therapeutic target for the mitochondrial DNA 3243A > G mutation-related disease. Cell Death Dis. 12, 701 (2021).

63. Geng, X. et al. Mitochondrial DNA mutation m.3243A>G is associated with altered mitochondrial function in peripheral blood mononuclear cells, with heteroplasmy levels and with clinical phenotypes. Diabet. Med. 36, 776–783 (2019).

64. Varanasi, S. S., Francis, R. M., Berger, C. E. M., Papiha, S. S. & Datta, H. K. Mitochondrial DNA Deletion Associated Oxidative Stress and Severe Male Osteoporosis. Osteoporosis Int 10, 143–149 (2019).

65. Catheline, S. E., Kaiser, E. & Eliseev, R. A. Mitochondrial Genetics and Function as Determinants of Bone Phenotype and Aging. Current osteoporosis reports 21, 540—551 (2023).

66. Stegen, S. & Carmeliet, G. Metabolic regulation of skeletal cell fate and function. Nat. Rev. Endocrinol. 1–15 (2024) doi:10.1038/s41574-024-00969-x.

67. Rowlands, V. et al. Optimisation of robust singleplex and multiplex droplet digital PCR assays for high confidence mutation detection in circulating tumour DNA. Sci. Rep. 9, 12620 (2019).

68. Fernandez-Guerra, P. et al. Bioenergetic and Proteomic Profiling of Immune Cells in Myalgic Encephalomyelitis/Chronic Fatigue Syndrome Patients: An Exploratory Study. Biomolecules 11, 961 (2021).

69. Morales, J. et al. A joint NCBI and EMBL-EBI transcript set for clinical genomics and research. Nature 604, 310–315 (2022).

70. Sahebekhtiari, N. et al. Deficiency of the mitochondrial sulfide regulator ETHE1 disturbs cell growth, glutathione level and causes proteome alterations outside mitochondria. BBA - Molecular Basis of Disease 1865, 126–135 (2019).

71. Walls, A. B., Bak, L. K., Sonnewald, U., Schousboe, A. & Waagepetersen, H. S. Brain Energy Metabolism. Neuromethods 73–105 (2014) doi:10.1007/978-1-4939-1059-5_4.

72. Biemann, K. & McCloskey, J. A. Application of Mass Spectrometry to Structure Problems. 1 VI. Nucleosides 2. *J*. Am. Chem. Soc. 84, 2005–2007 (1962).

73. Andersen, J. V., Skotte, N. H., Aldana, B. I., Nørremølle, A. & Waagepetersen, H. S. Enhanced cerebral branched-chain amino acid metabolism in R6/2 mouse model of Huntington’s disease. Cell. Mol. Life Sci. 76, 2449–2461 (2019).

74. Chen, L. & Ditzel, N. In vivo Heterotopic Bone Formation Assay Using Isolated Mouse and Human Mesenchymal Stem Cells. BIO-Protoc. 5, (2015).

75. Hansen, M. S. et al. GIP reduces osteoclast activity and improves osteoblast survival in primary human bone cells. Eur. J. Endocrinol. 188, 144–157 (2023).

76. Ding, M., Odgaard, A., Linde, F. & Hvid, I. Age-related variations in the microstructure of human tibial cancellous bone. J. Orthop. Res. 20, 615–621 (2002).

77. T, Hildebrand. & P, RÜegsegger. Quantification of Bone Microarchitecture with the Structure Model Index. Comput. Methods Biomech. Biomed. Eng. 1, 15–23 (1997).

78. Drejer, L. A. et al. Trabecular bone deterioration in a postmenopausal female suffering multiple spontaneous vertebral fractures due to a delayed denosumab injection – A post-treatment re-initiation bone biopsy-based case study. Bone Rep. 19, 101703 (2023).

79. Andersen, T. L. et al. A Physical Mechanism for Coupling Bone Resorption and Formation in Adult Human Bone. Am. J. Pathol. 174, 239–247 (2009).

80. Sørensen, N. N. et al. Disturbed bone marrow adiposity in patients with Cushing’s syndrome and glucocorticoid- and postmenopausal- induced osteoporosis. Front. Endocrinol. 14, 1232574 (2023).

81. Burt, V. L. & Harris, T. The Third National Health and Nutrition Examination Survey: Contributing Data on Aging and Health. 34, 486–490 (1994).

82. Hansen, M. S. et al. Transcriptional reprogramming during human osteoclast differentiation identifies regulators of osteoclast activity. Bone Res. 12, 5 (2024).

83. Love, M. I., Huber, W. & Anders, S. Moderated estimation of fold change and dispersion for RNA-seq data with DESeq2. Genome Biol. 15, 550 (2014).

84. Young, M. D., Wakefield, M. J., Smyth, G. K. & Oshlack, A. Gene ontology analysis for RNA-seq: accounting for selection bias. Genome Biol. 11, R14 (2010).

85. Kuleshov, M. V. et al. Enrichr: a comprehensive gene set enrichment analysis web server 2016 update. Nucleic Acids Res. 44, W90–W97 (2016).

86. Chen, E. Y. et al. Enrichr: interactive and collaborative HTML5 gene list enrichment analysis tool. BMC Bioinform. 14, 128 (2013).

