## Supplemental data for "Restoring oxidative phosphorylation enhances osteogenesis in mitochondrial DNA translation defective human bone marrow stromal cells"

### Supplementary tables

#### Table S1. Clinical, biochemical and heteroplasmy (m.3243A>G/wild-type) data.

| *Parameter* | **Controls (n=10)** | **m.3243A>G carriers (n=10)** | **Fold Change (m.3243A>G/Control)** | ***p*-value** |
|---|---|---|---|---|
| *Male/Female* | 4/6 | 4/6 | | |
| *Age (years)* | 48.0 (29 - 65) | 50.5 (26 - 58) | 1,05 | 0.109 |
| *Smokers* | 0 | 1 | | |
| *Postmenopausal females (age (years))* | 3 (46-50) | 4 (38-50) | | |
| *Hearing impairment* | - | 7/10 | - | - |
| *Diabetes* | - | 7/10 | - | - |
| *Myopathy* | - | 5/10 | - | - |
| *BMI (kg/m²)* | 23.6 (21.76 - 26.4) | 22.66 (17.21 - 29.32) | 0,96 | 0.77 |
| *Diabetes duration (years)* | 0 | 9 (0 - 39) | - | **0.021 *** |
| *Fasting glucose (mmol/L)* | 5.45 (4.8 - 6.3) | 6.75 (4.8 - 18.5) | 1,24 | 0.078 |
| *HbA1c (mmol/L)* | 36 (29 - 38) | 49.5 (31 - 80) | 1,38 | **0.02 *** |
| *Insulin (mlU/L)* | 54.5 (32 - 81) | 60.5 (17 - 297) | 1,11 | 0.432 |
| *C- peptide (pmol/L)* | 709 (452 - 921) | 650 (192 - 1888) | 0,92 | 0.77 |
| *Hb (mmol/L)* | 8.95 (8.2 - 10.1) | 8.4 (8.2 - 10.7) | 0,94 | 0.44 |
| *Trombocytes (10E9/L)* | 264.5 (208 - 293) | 258 (219 - 302) | 0.98 | 0.722 |
| *AST (U/L)* | 26.5 (20 - 47) | 33 (20 - 49) | 1,25 | 0.205 |
| *Creatinine (μmol/L)* | 79.5 (60 - 117) | 74 (49 - 116) | 0,93 | 0.155 |
| *Creatinine kinase (U/L)* | 96.5 (40 - 318) | 139 (73 - 545) | 1,44 | 0.375 |

| | | | | |
|---|---|---|---|---|
| *pH* | 7.37 (7.32 - 7.39) | 7.37 (7.21 - 7.47) | 1 | 0.343 |
| *Lactate (mmol/L)* | 1.3 (1 - 2.5) | 2.05 (1.6 - 5.5) | 1,58 | **0.006 **** |
| *Pyruvate (µmol/L)* | 143.5 (115 - 207) | 212.5 (80 - 310) | 1.48 | 0.058 |
| *Lactate/Pyruvate* | 9.11 (6.28 - 14.29) | 10.36 (9.13 - 20) | 1.14 | 0.193 |
| *Ammonium (µmol/L)* | 23 (13 - 38) | 22 (13 - 42) | 0,96 | 1 |
| *Sodium (mmol/L)* | 140 (138 - 146) | 137 (134 - 139) | 0.98 | **0.014 *** |
| *Potassium (mmol/L)* | 3.9 (3.6 - 4.6) | 4.1 (3.7 - 4.7) | 1.05 | 0.183 |
| *PTH (pmol/L)* | 4.75 (2.9 - 8.9) | 5.7 (2.2 - 8.8) | 1.2 | 0.541 |
| *ALP (U/L)* | 72.5 (53 - 96) | 89.5 (64 - 117) | 1,23 | 0.105 |
| *Calcium ion (mmol/L)* | 1.26 (1.2 - 1.37) | 1.25 (1.13 - 1.3) | 0,99 | 0.305 |
| *1.25-OH vitamin D3* | 63 (53 - 106) | 85 (42 - 125) | 1.35 | 0.25 |
| *TSH (10E-3IU/L)* | 1.4 (0.74 - 2.2) | 1.7 (0.82 - 3.6) | 1.21 | 0.16 |
| *eGFR/1.73m2* | 6.1 (5.1 - 6.4) | 8.05 (5.3 - 12.5) | 1,32 | **0.02 *** |
| *Heteroplasmy MSC (%)* | 0.01 (0 - 0.11) | 31.51 (12.51 - 81.92) | 3151 | **0.008 **** |
| *Heteroplasmy OC (%)* | 0 (0 - 0.16) | 21.94 (6.7 - 43.78) | - | **0.016 *** |

Data represents median (minimum – maximum). Significant *p*-values are highlighted by asterisk and calculated using Wilcox signed-ranked test.

ALP: alkaline phosphatase; AST: aspartate aminotransferase; BMI: body mass index; eGFR: estimated glomerular filtration rate; HbA1C: glycated haemoglobin; MSC: mesenchymal stem cells; OC: osteoclast; PTH: parathyroid hormone; TSH: thyroid stimulating hormone.

#### Table S2. Bioenergetic profile of hBMSC

| Parameter | Control | m.3243A>G | p-value | significance |
|---|---|---|---|---|
| ATP rate index | 0.73 (0.45 - 1.75) | 0.44 (0.09 - 1.12) | 0.002 | ** |
| ATP-linked respiration | 1.31 (0.81 - 1.93) | 1.05 (0.15 - 1.56) | 0.160 | ns |
| ATP-linked to maximal respiration | 0.38 (0.25 - 0.52) | 0.35 (0.18 - 0.45) | 0.064 | ns |
| Basal glycolysis | 10.69 (5.55 - 15.38) | 16.78 (9.94 - 20.32) | 0.002 | ** |
| Basal mitochondrial respiration / Basal glycolysis | 0.12 (0.06 - 0.23) | 0.08 (0.03 - 0.17) | 0.017 | • |
| Basal respiration | 1.36 (0.93 - 1.89) | 1.23 (0.39 - 1.57) | 0.160 | ns |
| Cell respiratory control | 11.98 (8.92 - 43.76) | 10.94 (2.58 - 13.73) | 0.084 | ns |
| Compensatory glycolysis | 15.77 (10.72 - 23.28) | 19.95 (14.07 - 28.24) | 0.105 | ns |
| Coupling efficiency | 92.84 (86.5 - 110.33) | 84.35 (37.53 - 99.64) | 0.105 | ns |
| Glycolyis (%) | 93.5 (87.86 - 96.62) | 95.53 (90.83 - 98.37) | 0.014 | • |
| Glycolytic ATP (%) | 58.28 (38.19 - 69.43) | 69.66 (47.49 - 91.99) | 0.002 | ** |
| Glycolytic ATP production | 9.84 (5.69 - 13.91) | 14.6 (7.59 - 20.05) | 0.009 | ** |
| Maximal respiration | 3.5 (3.05 - 4.02) | 3.12 (0.82 - 3.52) | 0.105 | ns |
| Mitochondrial ATP (%) | 41.72 (30.57 - 61.81) | 30.34 (8.01 - 52.51) | 0.002 | ** |
| Mitochondrial ATP production | 7.5 (4.74 - 10.84) | 6.14 (1.6 - 9.03) | 0.193 | ns |
| Non-glycolytic acidification | 4.16 (2.69 - 5.94) | 5.12 (3.83 - 7.41) | 0.064 | ns |
| Non-mitochondrial respiration | 0.9 (0.52 - 1.1) | 0.88 (0.5 - 1.08) | 0.375 | ns |
| Proton Leak | 0.28 (0.08 - 0.4) | 0.3 (0.22 - 0.35) | 0.492 | ns |
| Spare respiratory capacity | 1.8 (1.74 - 2.09) | 1.86 (1.44 - 2.13) | 0.275 | ns |
| Total acidification | 11.36 (6.33 - 16.24) | 17.43 (10.79 - 21.22) | 0.002 | ** |
| Total ATP production | 16.27 (14.29 - 21.09) | 19.62 (15.43 - 27.44) | 0.232 | ns |

Data is expressed as median and range. Analyzed using Wilcox signed-ranked test *: *p*-value<0.05, **: *p*-value <0.01 n = 20 (10 carriers and 10 controls matched by age, sex, and BMI). Bioenergetic analysis of hBMSC from each matched pair of m.3243A<G carriers and controls was done simultaneously in one Seahorse experiment with 6-8 technical replicates per parameter.

ATP: adenosine triphosphate

### Table S3. Bioenergetic profile of mature osteoclasts

| Parameter | Control | m.3243A>G | p-value |
|---|---|---|---|
| ATP rate index | 2.95 (0.49 - 6.21) | 2.04 (0 - 4.83) | 0.625 |
| ATP-linked respiration | 75.46 (50.66 - 136.43) | 71.93 (54.88 - 99.72) | 0.846 |
| ATP-linked to maximal respiration | 0.29 (0.15 - 0.36) | 0.25 (0.16 - 0.34) | 0.322 |
| Basal glycolysis | 16.61 (1.9 - 60.68) | 15.35 (2.18 - 52.03) | 0.922 |
| Basal respiration | 85.78 (64.88 - 152.91) | 84.32 (62.75 - 127.53) | 0.922 |
| Cell respiratory control | 13.44 (7.35 - 19.44) | 13.92 (9.24 - 18.86) | 1.000 |
| Coupling efficiency | 85.62 (75.66 - 92.24) | 86.45 (74.25 - 94.28) | 0.770 |
| Glycolytic ATP (%) | 25.54 (14.42 - 68.24) | 31.12 (17.45 - 53.94) | 0.910 |
| Glycolytic ATP production | 184.26 (51.94 - 692.76) | 165.14 (1.4 - 478.57) | 0.557 |
| Glycolytic spare capacity | 1.08 (0.92 - 1.64) | 1.07 (0.96 - 1.76) | 1.000 |
| Maximal glycolytic capacity | 19.54 (10.13 - 74.59) | 19.19 (7.77 - 61.74) | 0.770 |
| Maximal respiration | 306.64 (158.23 - 413.35) | 298.42 (162.27 - 410.22) | 0.846 |
| Mitochondrial ATP (%) | 74.47 (31.76 - 85.58) | 68.88 (46.06 - 82.55) | 0.910 |
| Mitochondrial ATP production | 432.53 (304.67 - 762.32) | 428.98 (3.02 - 622.68) | 0.557 |
| Non-glycolytic acidification | 10.46 (5.44 - 18.27) | 8.35 (5.2 - 13.06) | 0.275 |
| Non-mitochondrial respiration | 42.68 (27.01 - 60.69) | 39.39 (29.96 - 49.7) | 0.770 |
| Proton Leak | 20.29 (13.53 - 37.45) | 22.05 (13.88 - 34.53) | 1.000 |
| Spare respiratory capacity | 2.56 (1.9 - 3.82) | 2.66 (2.08 - 3.71) | 0.695 |
| Total ATP production | 715.7 (356.61 - 1029.53) | 613.98 (4.42 - 886.62) | 0.322 |

Data is expressed as median and range. Analyzed using Wilcox signed-ranked test *: $p$-value<0.05, **: $p$-value <0.01 n = 20 (10 carriers and 10 controls matched by age, sex, and BMI). Bioenergetic analysis of mature OCs (day 9 of induction) from each matched pair of m.3243A<G carriers (6-8 wells/parameter)

ATP: adenosine triphosphate; OC: osteoclast

### Table S4. Participants DXA scans

| | Controls (n=10) | m.3243A>G carriers (n=10) | *p*-value |
|---|---|---|---|
| Lumbar spine, total area ($mm^2$) | 59.6 ± 15.9 | 54.7 ± 9.4 | $p$ = 0.14 |
| Lumbar spine, total BMC ($mg/cm^3$) | 58.8 ± 23.5 | 49.7 ± 13.6 | $p$ = 0.12 |
| Lumbar spine, total BMD (T-score) SD | 0.96 ± 0.15 | 0.90 ± 0.12 | $p$ = 0.51 |
| Femur neck, area ($mm^2$) | 5.4 ± 0.6 | 5.3 ± 0.4 | $p$ = 0.25 |
| Femur neck, BMC ($mg/cm^3$) | 4.2 ± 0.9 | 3.7 ± 1.1 | $p$ = 0.46 |
| Femur neck, BMD (T-score) SD | 0.76± 0.12 | 0.69 ± 0.17 | $p$ = 0.30 |
| Hip, total areal ($mm^2$) | 38.3 [33.6-43.1] | 36.1 [31.3-41.7] | $p$ = 0.45 |
| Hip, total BMC ($mg/cm^3$) | 37.6 ± 12.9 | 29.7 ± 10.9 | $p$ = 0.62 |
| Hip, total BMD (T-score) SD | 0.93 ± 0.12 | 0.81 ± 0.19 | $p$ = 0.21 |

Bone areal, bone mineral content, bone density and T-scores by DXA in m.3243A>G carriers and controls.

Distribution of data was examined using the Shapiro-Wilk normality test. Data presented as mean +/- SD, or median [IQR] according to distribution. Paired test for parametric data and Wilcox signed-rank test for non-parametric data.

n = 20 (10 carriers and 10 controls matched by age, sex, and BMI).

BMC: bone mineral content; BMD: bone mineral density

### Supplementary figures

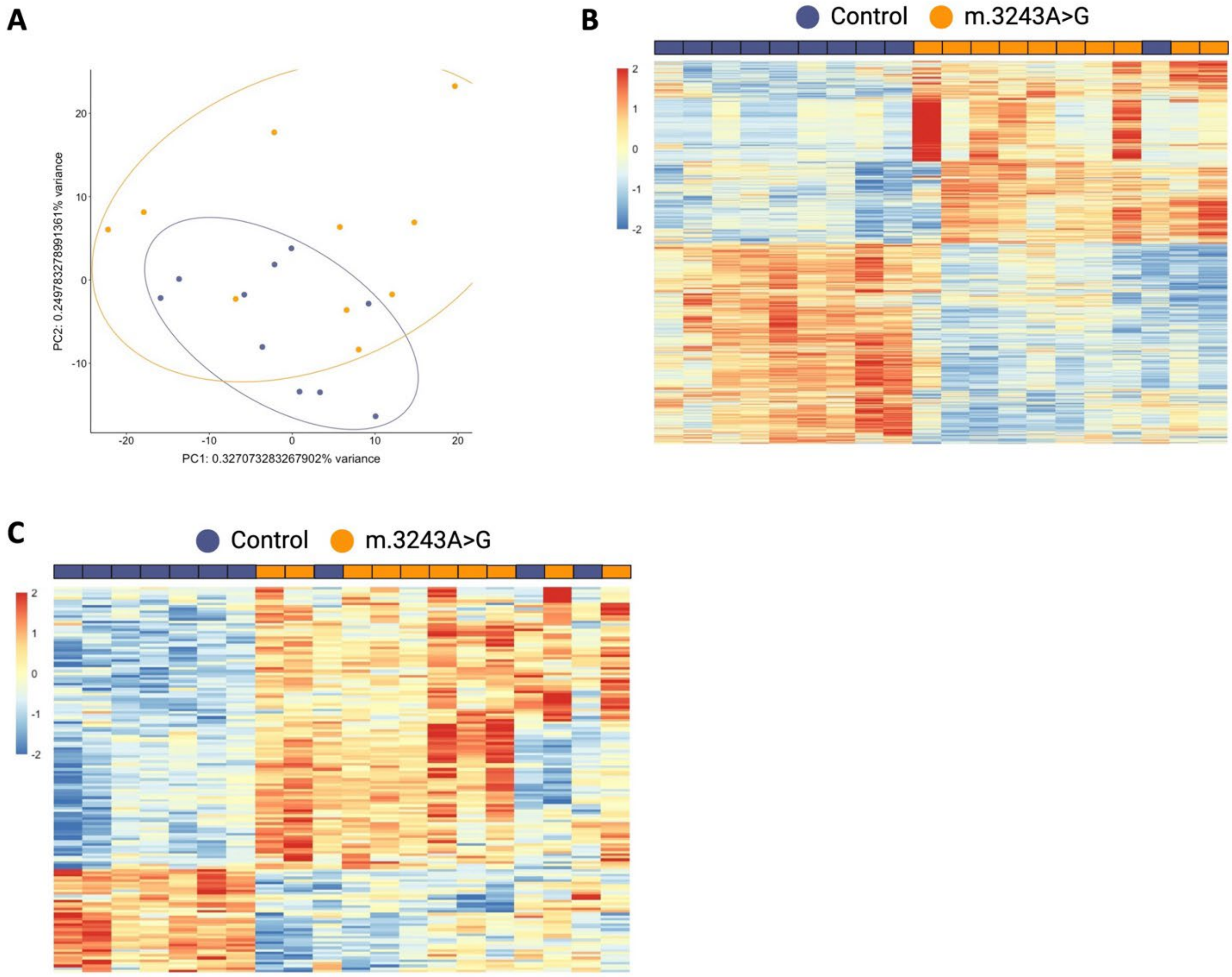

#### Figure S1. hBMSC from m.3243A>G transcriptional profile.

**A)** Unsupervised PCA of transcriptomic data from RNA sequencing (19,199 genes).

**B)** Heat map showing the fold-change (carriers/controls) for the 1,038 DE genes analysed by RNA sequencing.

**C)** Heat map showing the fold-change (carriers/controls) for the 149 mitochondrial genes DE in carriers analysed by RNA sequencing.

n = 20 (10 m.3243A>G carriers and 10 controls matched by age, sex, and BMI). RNA sequencing analysis was done in the same batch.

DE: differentially expressed; hBMSC: human bone marrow mesenchymal stem cell; PCA: principal component analysis

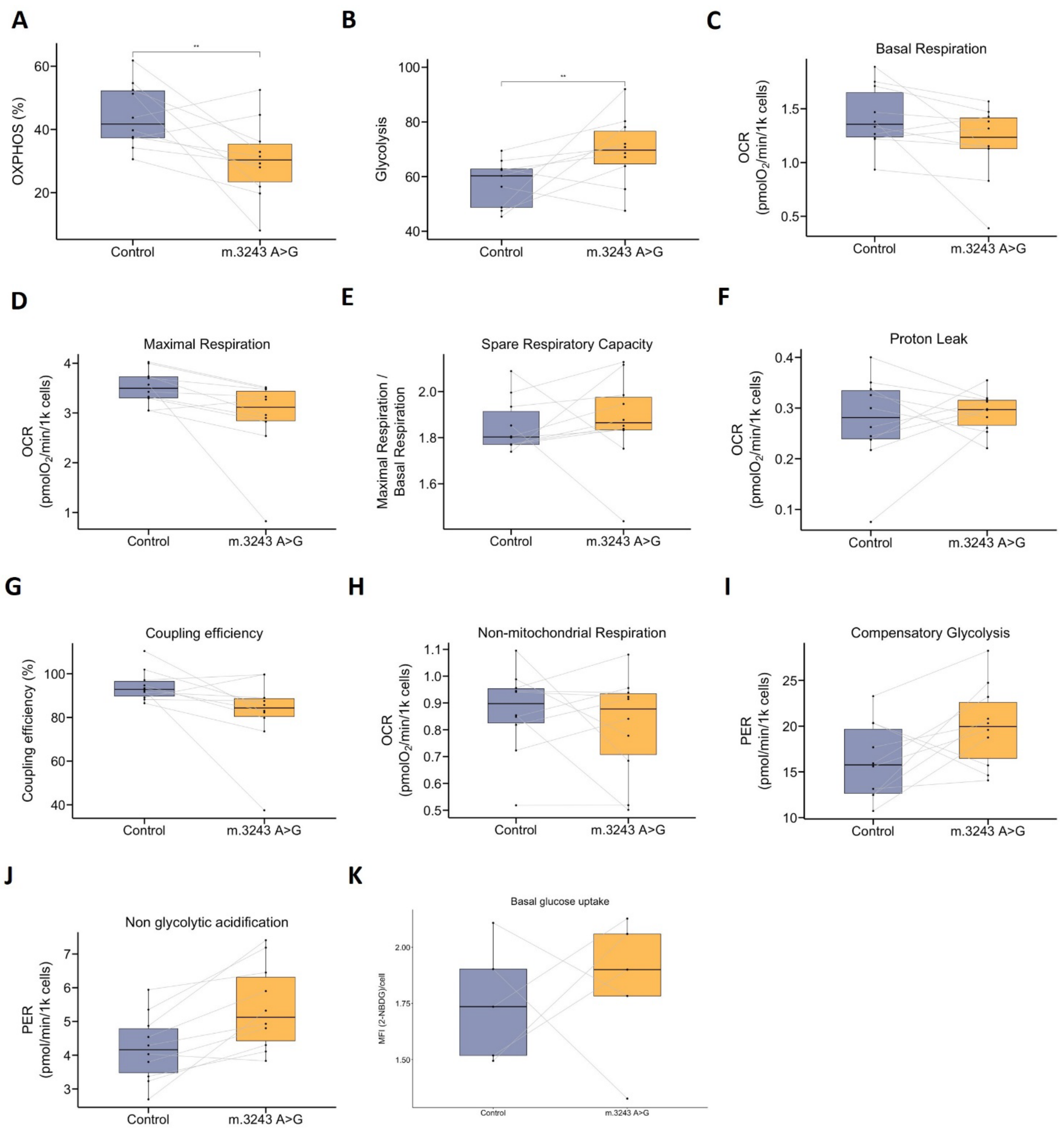

#### Figure S2. Bioenergetic profile of hBMSC from m.3243A>G carriers compared to controls.

**A-J)** Bioenergetics of hBMSC were analysed by extracellular flux analysis using Seahorse analyzer. The data was normalised to the number of Hoechst positive cells in each well analysed with Cytation1. n = 20 (10 carriers and 10 controls matched by age, sex, and BMI) (6-8 wells/group). **A)** Mitochondrial ATP production, **B)** Glycolytic ATP production, **C)** Basal respiration, **D)** Maximal respiration, **E)** Spare respiratory capacity, **F)** Proton leak, **G)** Coupling efficiency, **H)** Non-mitochondrial respiration, **I)** Compensatory glycolysis, **J)** Non-glycolytic acidification.

**K)** The box plots show the basal glucose uptake of hBMSC analysed by accumulation of the 2-NBDG fluorescent probe using Cytation1 high-content fluorescent microscope. n = 10 (5 carriers and 5 controls) (6 wells/group).

Bioenergetic analysis of matched m.3243A<G carriers and controls hBMSC was done in the same Seahorse plate. Glucose uptake of matched m.3243A<G carriers and controls hBMSC was done in the same plate. The data points of the box plot showed normalized data expressed as the mean of the technical replicate for each individual. Significance was calculated using Wilcox signed-ranked test *: *p*-value<0.05, **: *p*-value <0.01

ATP: adenosine triphosphate; 2-NBDG: 2-Deozy-[(7-nitro-2,1,3-benzoxadiazol-4-4yl)amino]-D-glucose; hBMSC: human bone marrow mesenchymal stem cell.

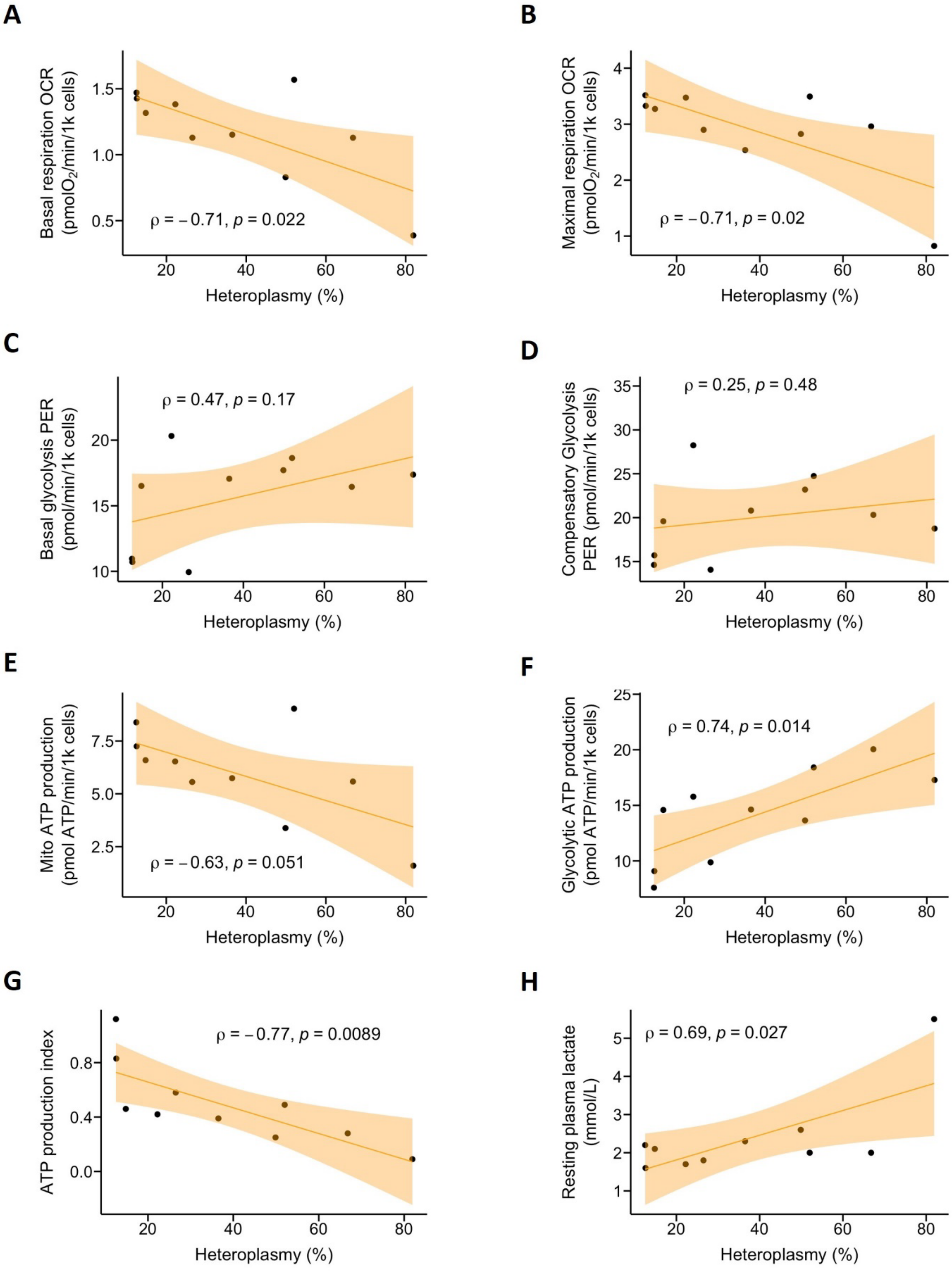
A
Basal respiration OCR (pmolO2/min/1k cells)
ρ = −0.71, p = 0.022
Heteroplasmy (%)
B
Maximal respiration OCR (pmolO2/min/1k cells)
ρ = −0.71, p = 0.02
Heteroplasmy (%)
C
Basal glycolysis PER (pmol/min/1k cells)
ρ = 0.47, p = 0.17
Heteroplasmy (%)
D
Compensatory Glycolysis PER (pmol/min/1k cells)
ρ = 0.25, p = 0.48
Heteroplasmy (%)
E
Mito ATP production (pmol ATP/min/1k cells)
ρ = −0.63, p = 0.051
Heteroplasmy (%)
F
Glycolytic ATP production (pmol ATP/min/1k cells)
ρ = 0.74, p = 0.014
Heteroplasmy (%)
G
ATP production index
ρ = −0.77, p = 0.0089
Heteroplasmy (%)
H
Resting plasma lactate (mmol/L)
ρ = 0.69, p = 0.027
Heteroplasmy (%)

#### Figure S3. Heteroplasmy correlation with bioenergetics parameters.

Spearman correlation between the percentage of heteroplasmy and

**A)** Basal respiration

**B)** Maximal respiration

**C)** Basal glycolysis

**D)** Compensatory glycolysis

**E)** Mitochondrial ATP production

**F)** Glycolytic ATP production

**G)** ATP production index

n = 20 (10 m.3243A>G carriers and 10 controls matched by age, sex, and BMI).

ATP: adenosine triphosphate; OCR: oxygen consumption rate; ECAR: extracellular acidification rate; PER: proton efflux rate

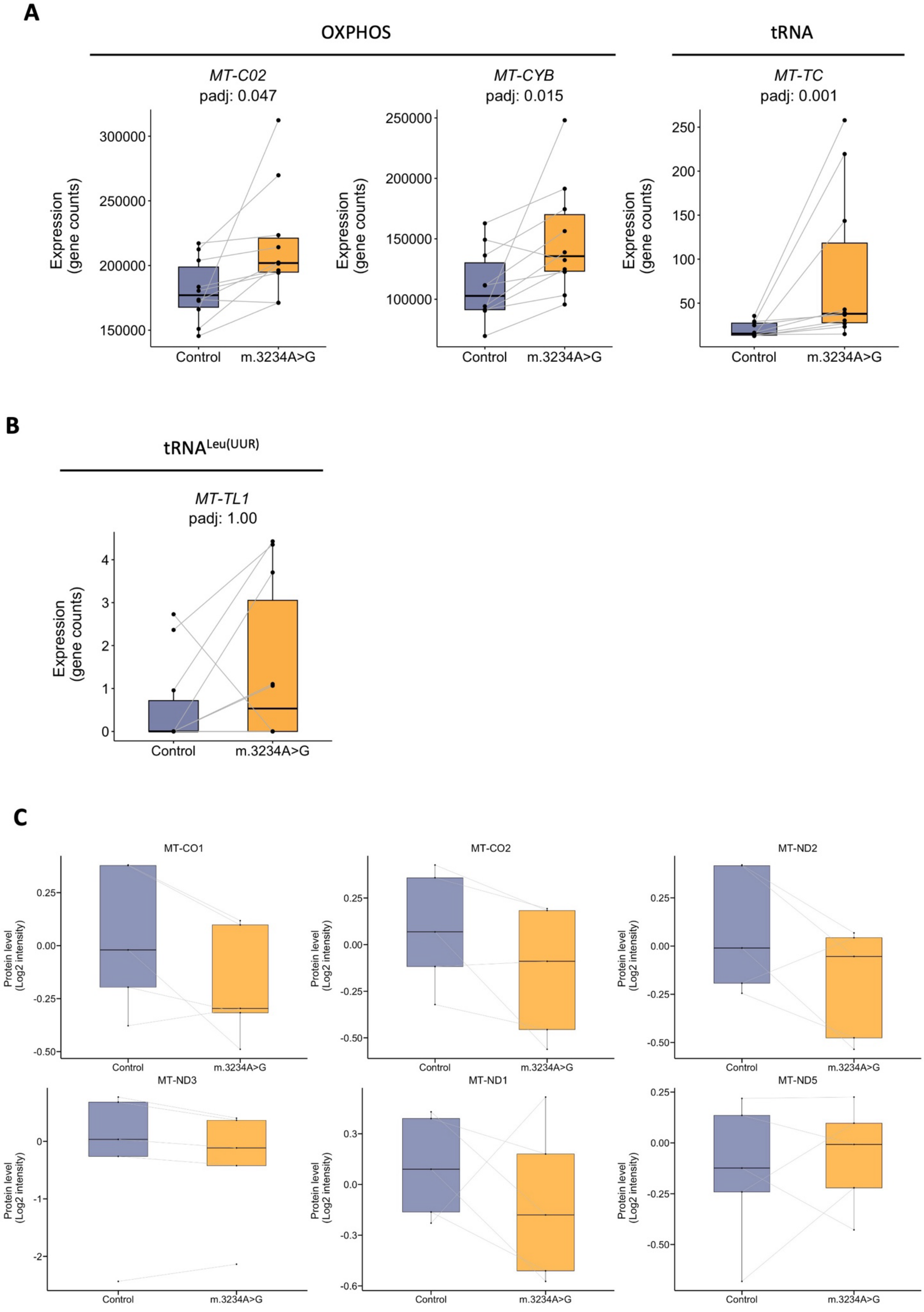
A
OXPHOS
tRNA
MT-C02
padj: 0.047
MT-CYB
padj: 0.015
MT-TC
padj: 0.001
Expression (gene counts)
Control
m.3234A>G
B
tRNALeu(UUR)
MT-TL1
padj: 1.00
C
MT-CO1
MT-CO2
MT-ND2
MT-ND3
MT-ND1
MT-ND5
Protein level (Log2 intensity)

#### Figure S4. mDNA encoded proteins.

**A)** Gene expression levels (read counts) of DE mDNA genes analysed by RNA sequencing

**B)** Gene expression levels of *MT-TL1* gene, where the pathogenic variant m.3243A>G is located

**C)** Boxplot of mDNA-encoded protein levels in hBMSC detected by mass spectrometry-based proteomics.

n = 20 (10 m.3243A>G carriers and 10 controls matched by age, sex, and BMI) for RNA sequencing. The statistical analysis was done using DESeq2 package. n = 10 (5 m.3243A>G carriers and 5 controls matched by age, sex, and BMI) for protein level analysis. Box plot showed normalised data expressed as each individual's mean of technical replicates. RNA sequencing and proteomics analysis were done in the same batch.

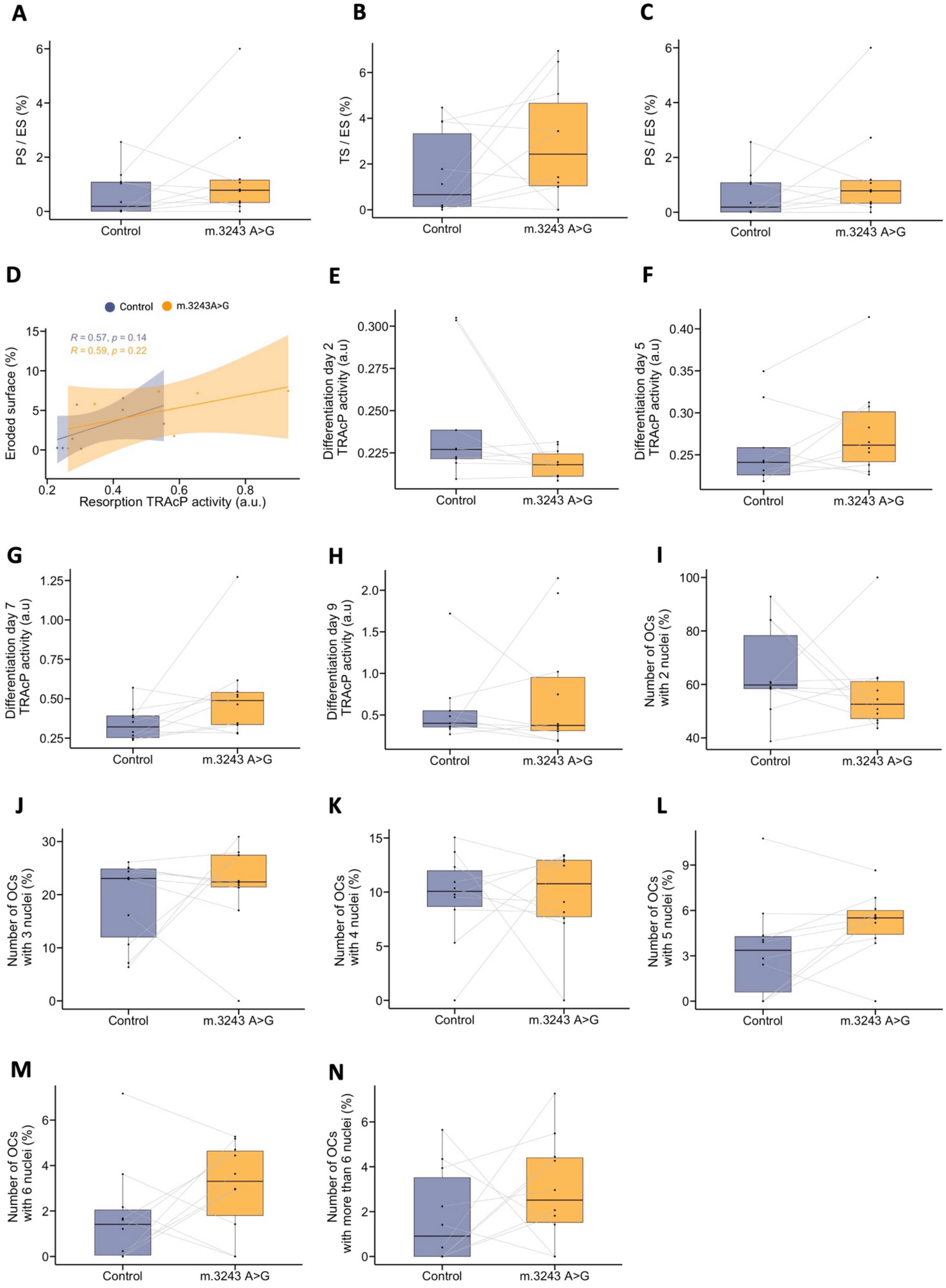
A
PS / ES (%)
Control
m.3243 A>G
B
TS / ES (%)
Control
m.3243 A>G
C
PS / ES (%)
Control
m.3243 A>G
D
Control
m.3243A>G
R = 0.57, p = 0.14
R = 0.59, p = 0.22
Eroded surface (%)
Resorption TRAcP activity (a.u.)
E
Differentiation day 2 TRAcP activity (a.u)
Control
m.3243 A>G
F
Differentiation day 5 TRAcP activity (a.u)
Control
m.3243 A>G
G
Differentiation day 7 TRAcP activity (a.u)
Control
m.3243 A>G
H
Differentiation day 9 TRAcP activity (a.u)
Control
m.3243 A>G
I
Number of OCs with 2 nuclei (%)
Control
m.3243 A>G
J
Number of OCs with 3 nuclei (%)
Control
m.3243 A>G
K
Number of OCs with 4 nuclei (%)
Control
m.3243 A>G
L
Number of OCs with 5 nuclei (%)
Control
m.3243 A>G
M
Number of OCs with 6 nuclei (%)
Control
m.3243 A>G
N
Number of OCs with more than 6 nuclei (%)
Control
m.3243 A>G

#### Figure S5. Osteoclasts characterization.

**A-C)** Resorptive activity of mature OCs after 3 days of culture in bone slices.

**A)** Pit-eroded surface quantified by eye after staining with touludin blue.

**B)** Trench-eroded surface quantified by eye after staining with touludin blue.

**C)** Ratio of pits- to trenches-eroded surface.

**D)** Pearson correlation between eroded surface and TRAcP activity released during resorption.

**E-H)** Extracellular TRAcP activity released during differentiation quantified by colorimetric substrate reduction.

**E)** TRAcP released during the first two days of differentiation.

**F)** TRAcP released from the second to the fifth day of differentiation.

**G)** TRAcP released from the fifth to the seventh day of differentiation.

**H)** TRAcP released from the seventh to the ninth day of differentiation.

**I-N)** Number of nuclei per mature OC quantified by brightfield imaging: with 2 nuclei **(I)**, with 3 nuclei **(J)**, with 4 nuclei **(K)**, with 5 nuclei **(L)**, with 6 nuclei **(M)**, with more than 6 nuclei **(N)**.

n = 20 (10 carriers and 10 controls matched by age, sex, and BMI). The statistical analysis performed was Wilcox signed-ranked test.

ES: eroded surface; OC: osteoclast; PS: pit surface; TS: trench surface; TRAcP: tartrate-resistanct acid phosphatase

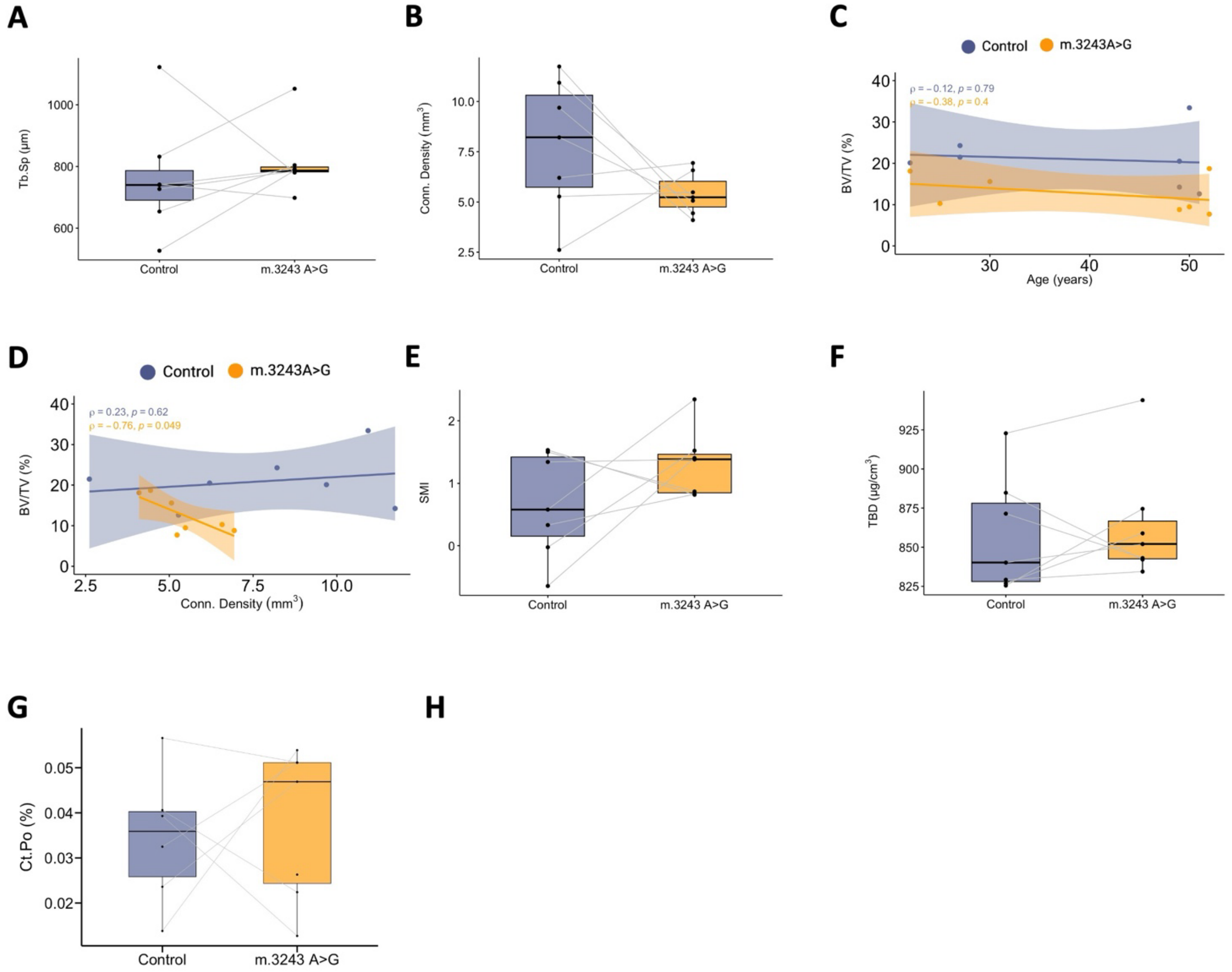

### Figure S6. Histomorphometry of transiliac crest biopsies.

**(A)** Trabecular separation (Tb.Sp) determined by µCT.

**(B)** Connective density determined by µCT.

**(C-D)** Spearman correlation analysis between BV/TV (%) and age **(C)** and Tb.Sp. **(D)**

**(E)** SMI.

**(F)** Trabecular bone density (TBD).

**(G)** Cortical porosity (Ct.Po).

n = 14 (7 carriers and 7 controls matched by age and sex). The statistical analysis performed was Wilcox signed-ranked test.

BV: bone volumen; Ct.Po: cortical porosity; SMI: structure model index; TBD: trabecular bone density; Tb.Sp: trabecular separation; TV: trabecular volumen;

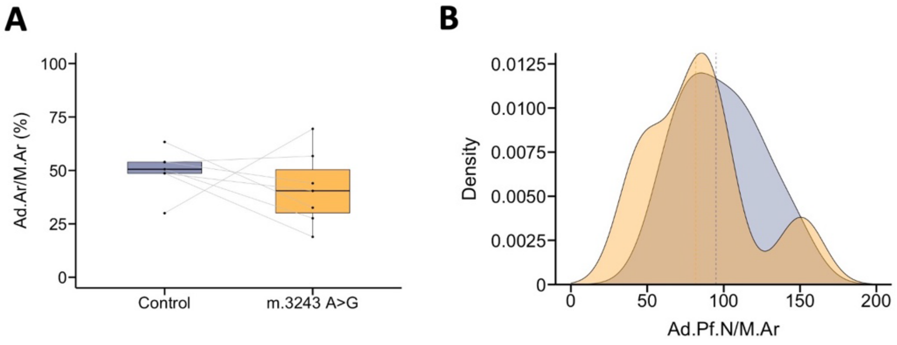

### Figure S7. Adipocyte histomorphometry

**A)** Adiposity area per marrow area (Ad.Ar/M.Ar)

**B)** Adipocyte profile number per marrow area (Ad.Pf.N/M.Ar)

n = 14 (7 carriers and 7 controls matched by age and sex). The statistical analysis performed was Wilcox signed-ranked test.

Ad.Ar/M.Ar: adiposity area per marrow area; Ad.Pf.Dm: adipocyte profile diameter; Ad.Pf.N/M.Ar: Adipocyte profile number per marrow area

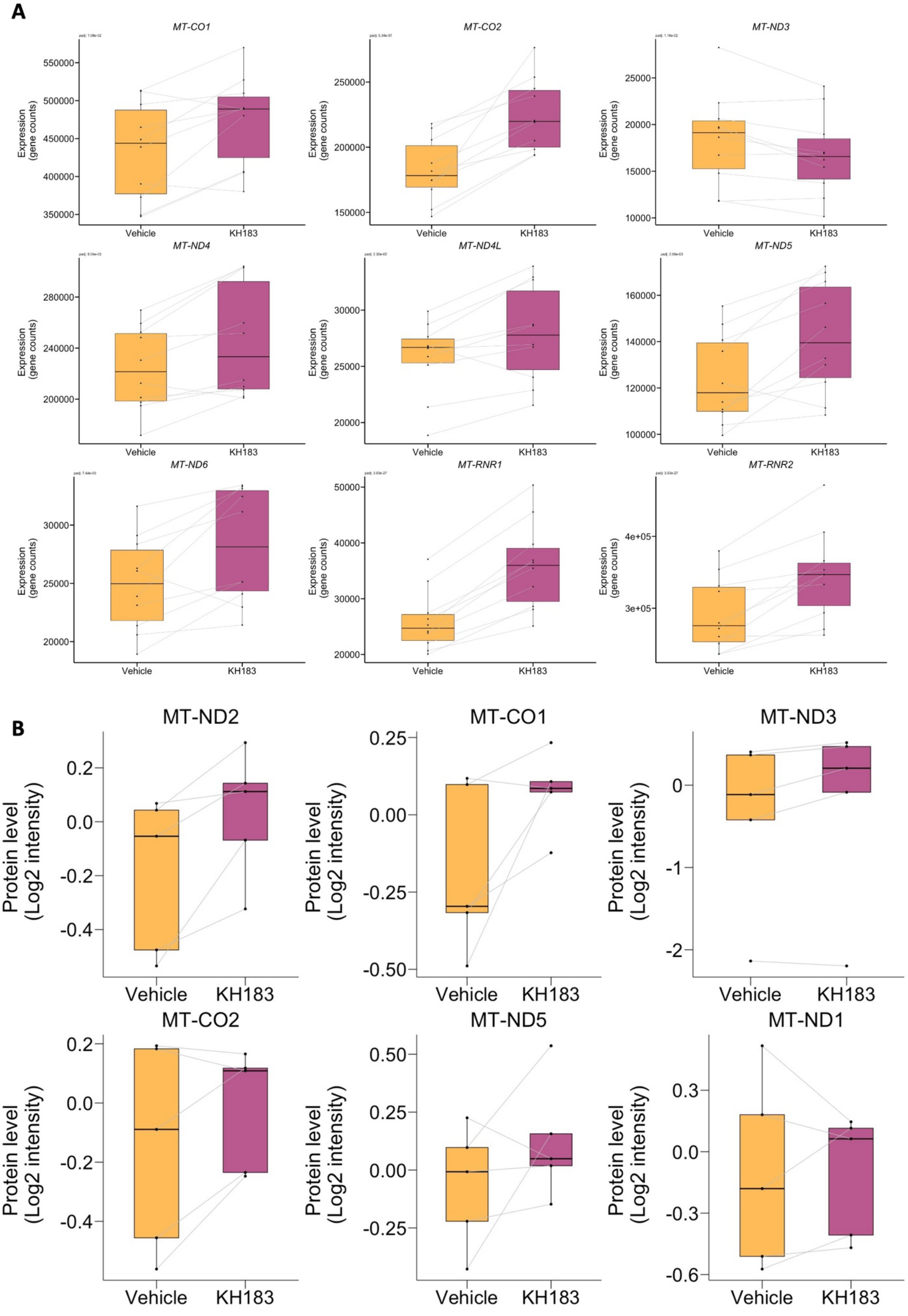
A
MT-CO1
MT-CO2
MT-ND3
MT-ND4
MT-ND4L
MT-ND5
MT-ND6
MT-RNR1
MT-RNR2
Expression (gene counts)
Vehicle
KH183
B
MT-ND2
MT-CO1
MT-ND3
MT-CO2
MT-ND5
MT-ND1
Protein level (Log2 intensity)
Vehicle
KH183

#### Figure S8. KH183 regulation of mDNA encoded genes

**A)** Boxplot of DE mDNA gene expression levels (read counts) in hBMSC analyzed by RNAsequencing

**B)** Boxplot of proteins encoded in DE mDNA genes in hBMSC analyzed by mass spectrometry-based proteomics.

n = 16 (8 m.3243A>G carriers´ hBMSC treated with KH183 or DMSO). Box plot showed normalised data expressed as each individual's mean of technical replicates. RNA sequencing and proteomics were done in the same batch. The statistical analysis performed was Wilcox signed-ranked test.

DE: differentially expressed; DMSO: Dimethyl sulfoxide

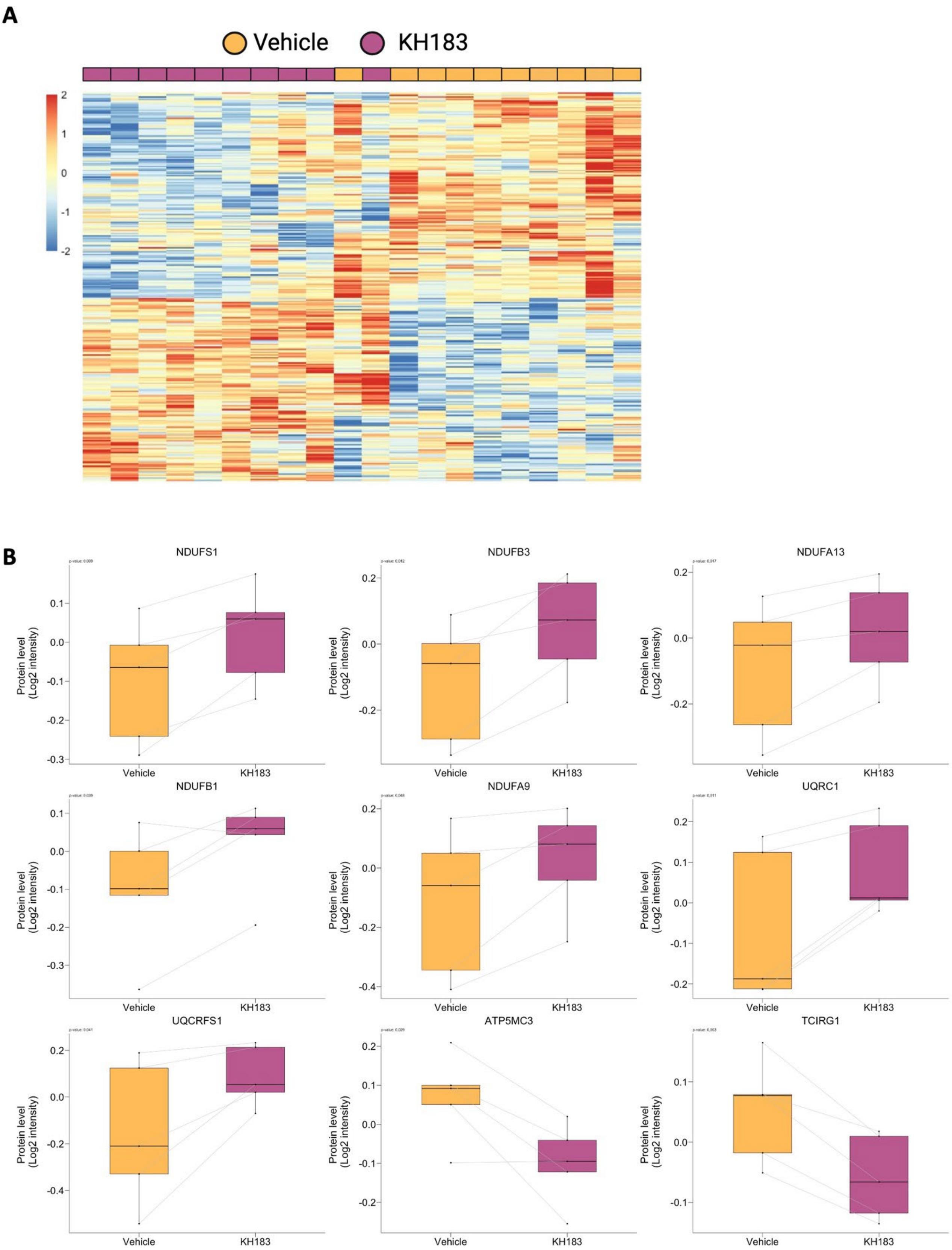
A
Vehicle
KH183
2
1
0
-1
-2
B
NDUFS1
NDUFB3
NDUFA13
NDUFB1
NDUFA9
UQRC1
UQCRFS1
ATP5MC3
TCIRG1
Protein level
(Log2 intensity)
Vehicle
KH183

#### Figure S9. KH183 regulation of OXPHOS genes

**A)** Heatmap of the number of DE genes encoding mitochondrial proteins (291 genes) in hBMSC from m.3243A>G after treatment with KH183 analysed by RNA sequencing

**B)** Boxplot of DE OXPHOS proteins levels in hBMSC from m.3243A>G after treatment with KH183 analysed by mass spectrometry-based proteomics.

n = 20 (10 m.3243A>G carriers´ hBMSC treated with KH183 or DMSO) for gene expression study. n = 10 (5 m.3243A>G carriers´ hBMSC treated with KH183 or DMSO) for protein level analysis

Box plot showed normalized data expressed as each individual's mean of technical replicates. RNA sequencing and proteomics were done in the same batch. The statistical analysis performed was Wilcox signed-ranked test.

DMSO: Dimethyl sulfoxide; DE: differentially expressed; OXPHOS: oxidative phosphorylation

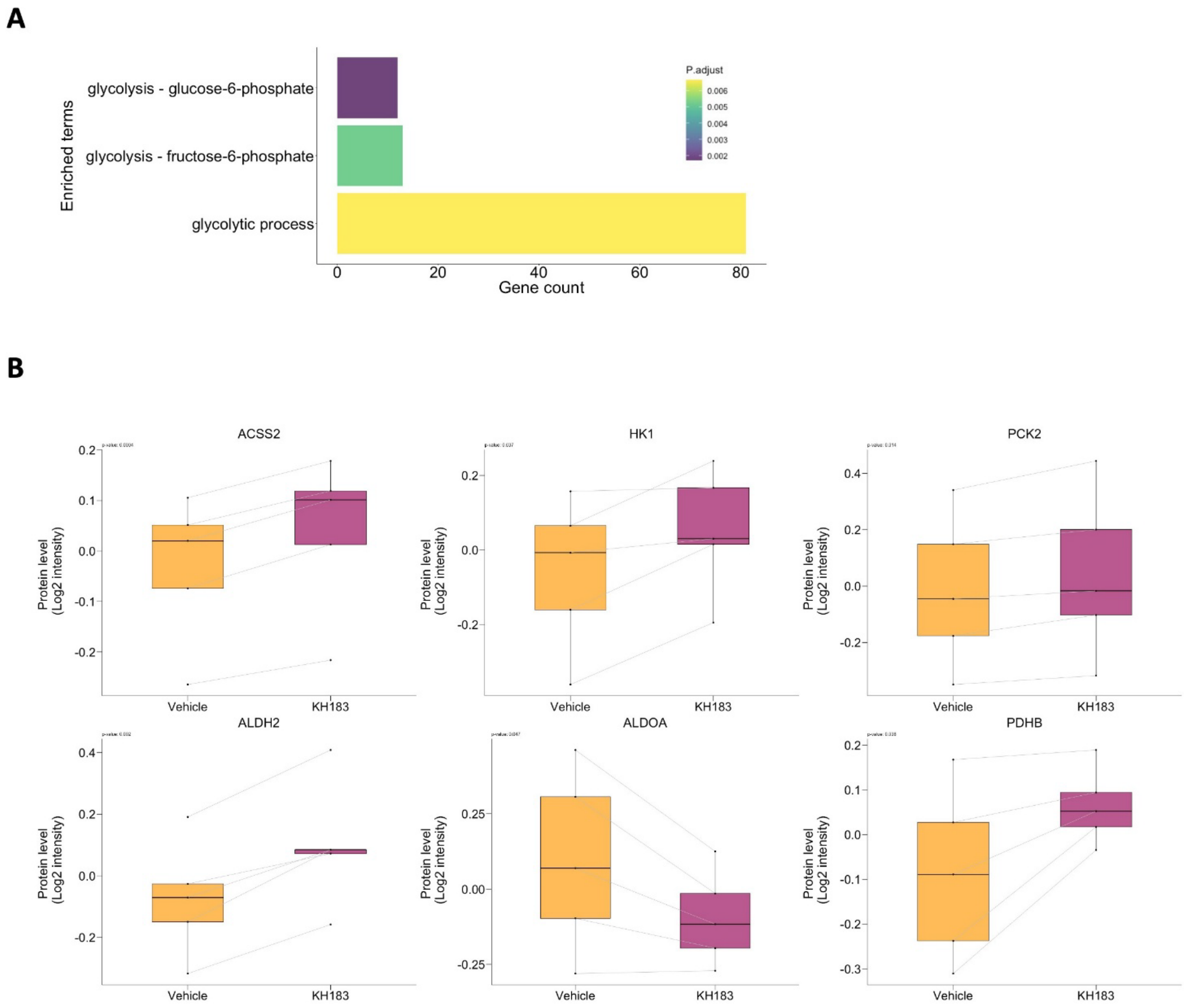

### Figure S10. KH183 regulation of glycolysis genes

**A)** Enrichment analysis of DE genes related to glycolysis in hBMSC from m.3243A>G after treatment with KH183 using GO biological process terms as database. The enriched terms show the number of genes in each GO term and the adjusted *p*-value of the GO term.

**B)** Boxplot of DE glycolysis proteins levels in hBMSC from m.3243A>G after treatment with KH183 analysed by mass spectrometry-based proteomics.

n = 20 (10 m.3243A>G carriers´ hBMSC treated with KH183 or DMSO) for gene expression study. n = 10 (5 m.3243A>G carriers´ hBMSC treated with KH183 or DMSO) for protein level analysis

Box plot showed normalised data expressed as each individual's mean of technical replicates. RNA sequencing and proteomics were done in the same batch. The statistical analysis performed was Wilcox signed-ranked test.

DMSO: Dimethyl sulfoxide
